# *Plasmodium*, the *Apicomplexa* outlier when it comes to protein synthesis

**DOI:** 10.1101/2023.08.10.552757

**Authors:** José R. Jaramillo Ponce, Magali Frugier

## Abstract

*Plasmodium* is an obligate intracellular parasite that makes numerous interactions with different hosts during its elaborate life cycle. This is also the case for other parasites that belong to the same phylum *Apicomplexa*. In this study, we identified bioinformatically the components of the multi-synthetase complexes (MSC) of several *Apicomplexa* parasites. By using AlphaFold2 modeling to compare their assembly, it appears that none of these MSCs resemble those identified in *Plasmodium*. In particular, the discrepancies between the core components of *Plasmodium* complexes, tRip and its homologs indicate that tRip-dependent exogenous tRNA import is not conserved in the other *Apicomplexa* parasites. Based on this observation, we looked for obvious differences that could explain this singularity in *Plasmodium*. The content of tRNA genes and amino acid usage in the different genomes highlighted the originality of *Plasmodia* translation. This is evident with respect to asparagine amino acid, which is highly used in the *Plasmodium* proteomes, and the scarcity of tRNA^Asn^ required for protein synthesis, regardless of long homorepeats or AT content of the genomes.

## INTRODUCTION

*Apicomplexa* is a group of obligate intracellular parasites with more than 6000 described species (1). Many of these parasites are important pathogens of humans. The *Apicomplexa* phylum includes (i) *Plasmodium*, the parasite responsible for malaria, a mosquito-borne disease that is potentially deadly (2,3); (ii) *Toxoplasma gondii* is a source of toxoplasmosis associated to congenital neurological birth defects (for example, encephalitis and ocular disease) (4,5); (iii) *Cryptosporidia* cause opportunistic infections in immunosuppressed patients transmitted *via* contaminated food or water (6,7). Further, numerous infectious diseases in wild and domesticated animals are caused by members of the apicomplexan genera Babesia, Theileria, and Neospora.

Apicomplexan parasites have an unusual biology as intracellular eukaryotes growing inside another eukaryotic cell, which differentiates them from other pathogens. One of the most fascinating aspects that defines the biology of *Apicomplexa* is their ability to manipulate their host cells (8). During infection, the parasites cause changes not only in signaling or evasion of host immunity but also manipulate host cells to suit their own needs, such as supplying nutrients and molecular building blocks (amino acids, nucleotides, glucose…) to the parasite. Among the many strategies put in place to allow optimal development in the host cell, a singular example is provided by *Plasmodium* which is characterized by the presence of tRip (tRNA import protein), a unique tRNA transporter that also participates in the formation of the parasite multi-aminoacyl-tRNA synthetases complexes (MSC).

Aminoacyl-tRNA synthetases (aaRSs) are essential enzymes that ensure the attachment of a specific amino acid to its cognate tRNAs (9); In eukaryotes, a subset of cytosolic aaRSs are organized into MSC built on multifunctional aaRS-interacting proteins (AIMPs) (reviewed in (10–12). Despite their diversity, the assembly of these MSCs in protozoa follows a dominant strategy involving GST-like domains. Such domains are found to be essentially fused to AIMPs, aaRSs and elongation factors subunits and interact with each other *via* two well identified binding surfaces called interfaces 1 and 2. In *Plasmodium*, tRip is an AIMP like no other: It homodimerizes using an alternative interface, named 1’ (13,14), it is an integral membrane protein anchored in the plasma membrane of the parasite (16); It allows the formation of two MSCs characterized by two different GST-like domain organizations (14,16). Most importantly, tRip is involved in the import of exogenous (host) tRNAs into the parasite and its deletion leads to low infectivity and translational efficiency which especially affects asparagine-rich proteins (15,17).

In this study, we investigated the possibility for other parasites of the *Apicomplexa* phylum to form MSC equivalent to those found in *Plasmodium*. To reconstruct the different MSCs’ architectures, sequence analysis and AlphaFold2 modeling were combined. This work was performed on 4 other genera than *Plamodiidae* by selecting several parasites with complete and annotated genomes in *Babesiidae*, *Theileriidae*, *Cryptosporidiidae* and *Sarcocystidae* (Figure 1). In the absence of conserved MSC assemblies, we searched for translational constraints that make *Plasmodium* such an unconventional parasite.

**Figure 1.**
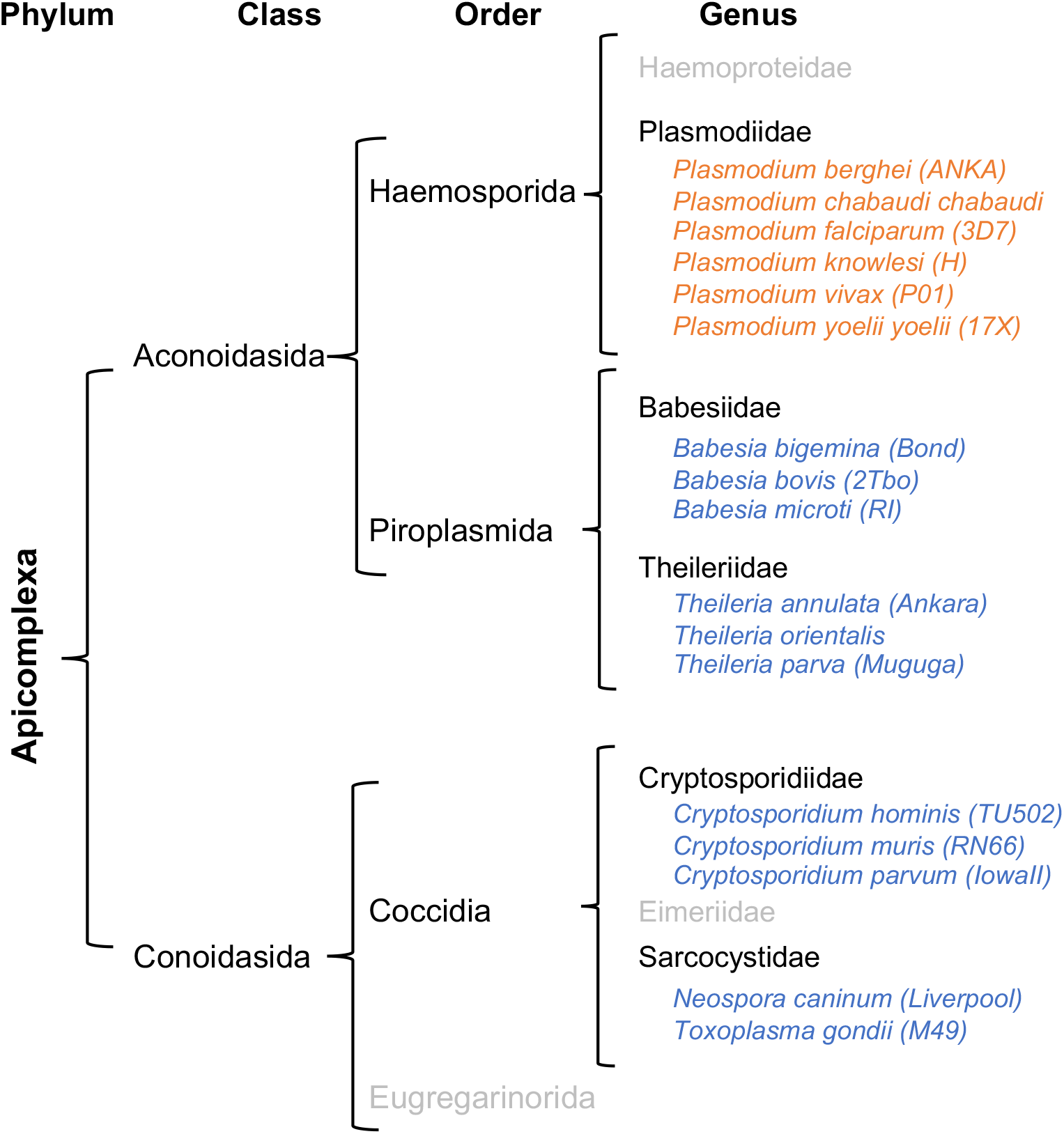
Phylogeny of *Apicomplexa* parasites. Most members of the *Apicomplexa* phylum are obligate parasites, some of which cause diseases in vertebrates. Their life cycle consists of three stages: sporozoite (infective stage), merozoite (asexual reproduction) and gametocyte (sexual reproduction). *Apicomplexa* are characterized by the presence of an apical complex responsible for the invasion of the parasite into the host cells and most of them possess an apicoplast, a plastid essential for their survival (reviewed in (44)). The phylum is divided into two classes: *Aconoidasida* and *Conoidasida*. On the one hand, the *Conoisada* include all species of *Cryptosporidium*: *C. hominis*, *C. parvum*, *C. muris*, among others, as well as *Neospora* and *Toxoplasma*. On the other hand, the *Aconoidasida* can be classified into *Haemosporida* consisting in *Plasmodium* species and *Piroplasmida* that include *Theileria* and *Babesia* species. This classification is the one used by the EupathDB database (45).

## MATERIAL AND METHODS

### Sequence retrieval and analysis

Entire proteomes, as well as individual sequences (proteins and tRNAs) were obtained directly from EuPathDB (Eukaryotic Pathogen DataBase) (18) (Figure S1). When protein genes were not annotated in the genomes, they were manually identified by Blast (19).Protein sequences alignments were performed with M-Coffee (20) to identify and delineate additional modules (GST-like, EMAPII-like, extra helices). Since GST-like domains have only been found in aaRSs and elongation factor 1 subunits to date, only these proteins have been investigated. As for tRNA sequences, when not annotated, they were identified using the tRNAScan-SE gene detection program (21) on *B. bigemina*, *T. annulata* and *C. parvum* genomes. Eventually, tRNA sets were completed by Blast. Box plots and line plots were generated with Excel.

### Complex modeling with AlphaFold2

Protein–protein complex predictions were generated in ColabFold (22) using https://colab.research.google.com/github/sokrypton/ColabFold/blob/main/AlphaFold2.ipynb

(Figure S1). Input protein sequences are those used for protein sequence alignments (Figure S2). Figures of structures were analyzed and generated using PyMOL version 2.1 (Schrodinger, Inc.). Only complexes with predicted template modeling scores (pTM) higher than 0.5 and showing potential interfaces 1, 1’ and 2 were considered (Figures S4 and S6). AlphaFold2 confidence estimates are shown for all models presented in Figures 3, 4, S3 and S5 (Figure S4, S6 and S7). Individual GST-like domains, all combinations of dimers (Figure 3A) and the oligomeric GST-like backbones of MSCs (Figure 3B) were predicted using the “no template information” option in ColabFold. For *Babesia bovis*, modeling of the GST-like backbone was performed with custom templates of *Bb*-p43, ERS and MRS obtained from predictions of heterodimers showing interfaces 1 and 2. For *Plasmodium berghei*, the Q- and M-complexes were predicted using the crystal structures *P. vivax* tRip (5ZKE), *P. berghei* ERS (8BCQ) and Raptor X models of QRS et MRS as custom templates (14). For *Toxoplasma gondii*, dimeric GST-like heterotrimers were predicted using Raptor X models of *Tg*-p43, ERS, QRS and MRS, all of them showing the expected GST-like fold and consistent with ColabFold predictions.

## RESULTS

### Identification of GST-like domains

The sequences of all cytosolic aaRSs and of elongation factor 1 (EF1) subunits were retrieved from the nuclear genomes of the selected apicomplexan parasites (Figure 1). A set of 20 cytosolic aaRSs was identified in all parasites, 4 EF1 subunits in *Plasmodiidae* and *Sarcocystidae* (EF1α, β, γ, δ, respectively in EupathDB, corresponding to EF1A, EF1Bα, EF1B³, EF1Bβ), and only 3 (EF1α, β, γ) in *Babesiidae*, *Theileriidae* and *Cryptosporidiidae* (Table S1). Proteins were aligned to identify GST-like domains potentially involved in complex formation and their structure was verified by ColabFold modeling (Figure S1). As expected, putative GST-like domains were found at the N-terminus of all EF1β, EF1γ, AIMPs and some of the ERSs, MRSs and QRSs (Table 1). They consist of two subdomains: the N-terminal thioredoxin-fold (β1-α1-β2-β3-β4-α2) and the C-terminal subdomain (α3 to α7) (Figure 2A); In the C-terminal domain, α-helices (α3 to α7) organize around the central helix α5, all are parallel one to each other except for α7. Helix α5 is mostly composed of hydrophobic residues and exhibits the N-capping box and hydrophobic staple motif (φ-S/T-X-X-D-φ), which is important for the stability of the fold (23) (Figure S2). However, sequence alignments and structure analysis showed that the N-capping box is missing (i) in all *Apicomplexa* MRS sequence but that of *Plasmodia* (Figure S2E). Several GST-like domains contained additional structures, specifically (i) the helix α8 at the C-terminus of EF1γ, ERS, MRS and some AIMPs, (ii) the insertion of one (EF1-γ, *Plasmodiidae* MRS and *Toxoplasma gondii* p43) or more α-helices (*Babesiidae*, *Theileriidae* and *Sarcocystidae* MRS) in the loop between strands β2 and β3 and (iii) an N-terminal extension in *T. gondii* ERS (Figure S2D). The GST-like domains of EF1β are exceptions as they do not contain a thioredoxin domain and are downsized to only 3 helices including α5 and α7 (Figure 2A).

**Figure 2.**
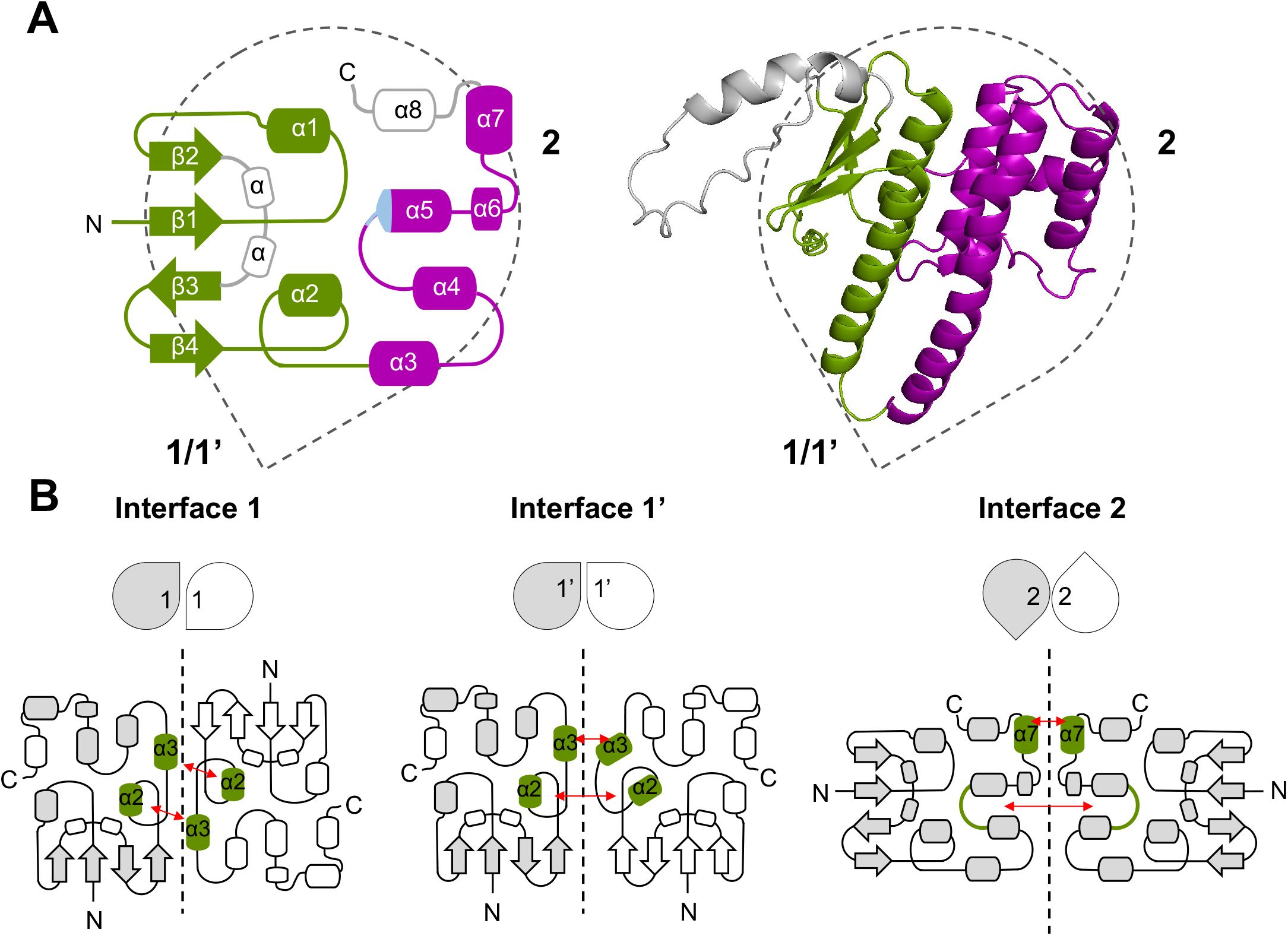
Structure and oligomerization of GST-like domains in eukaryotic MSCs and EF1. (**A**) Topological diagram and cartoon representation of a GST-like domain. The drop shape represents the orientation of the GST-like domain with oligomerization interfaces 1, 1’ and 2 highlighted. All secondary structures (α-helices and β-strands), and the N- and C-terminal ends are indicated. The thioredoxin-like subdomain (β1-α1-β2-β3-β4-α2) is colored in green, and the C-terminal helical subdomain is shown in purple (helices α3 to α7). Additional α-helices observed in some *Apicomplexa* GST-like domains are in grey. The position of the N-capping box and hydrophobic staple motif in helix α5 is colored in light blue. The model depicted in cartoon corresponds to the GST-like domain of *T. gondii* MRS predicted with ColabFold in complex with *Tg*-p43 and ERS. (**B**) Interaction interfaces involved in homo- and hetero-dimerization of GST-like domains. In each case, the drop shape representation, and the topological diagram of the GST-like dimer are shown. Interacting helices are colored in green and their contact pattern are indicated with red arrows. Interface 1 corresponds to a classical GST dimer, the two monomers being related by a 2-fold axis and interacting mainly through helices α2 and α3 in a parallel orientation. Interface 1’ is only observed in the crystal structure of *P. vivax* tRip, in which the N-termini of the two monomers are located on the same side of the homodimer and the interacting helices α2 and α3 are oriented perpendicularly. Interface 2 involves helix α7 and the loop connecting helices α4 and α5. A stacking interaction between two arginines from helices α7 is essential for dimerization and these residues are conserved only in GST-like domains from EF1-β, EF1-γ, ERS and AIMPs.

**Table 1.** Proteins potentially involved in GST-like driven complexes in *Apicomplexa* parasites. For each *Apicomplexa* genera, homologous proteins (blue) to *Plasmodium* GST-like-containing proteins (orange) were identified by Blast. Their additional domains located at their C-terminus are shown as well as the presence of a putative interface 2 in the GST-like domains (indicated by α 7).

|  | AIMPs and AARSs |  |  |  | Translation factors |  |
| --- | --- | --- | --- | --- | --- | --- |
|  | AIMP | ERS | MRS | QRS | EF1-β | EF1-γ |
| *Plasmodium* | GST (α7) | GST (α7) | GST | GST | GST (α7) | GST (α7) |
| *Babesia* | GST (α7) | GST (α7) | GST | - | GST (α7) | GST (α7) |
| *Theileria* | GST (α7) | GST (α7) | GST | - | GST (α7) | GST (α7) |
| *Cryptosporidium* | GST | - | - | - | GST (α7) | GST (α7) |
| *Toxoplasma* | GST (α7) | GST (α7) | GST | GST | GST (α7) | GST (α7) |
| *Neospora* | GST (α7) | GST (α7) | GST | GST | GST (α7) | GST (α7) |
| *Plasmodium* | EMAPII | - | EMAPII | K/R-rich helix |  |  |
| *Babesia* | EMAPII | - | - | K/R-rich helix |  |  |
| *Theileria* | EMAPII | - | - | helix bundle |  |  |
| *Cryptosporidium* | EMAPII | - | - | K/R-rich helix |  |  |
| *Toxoplasma* | EMAPII | - | EMAPII | K/R-rich helix |  |  |
| *Neospora* | EMAPII | - | EMAPII | K/R-rich helix |  |  |

### Description of GST-like interactions

Oligomerization of GST-like domains involves two canonical interfaces: Interface 1 and interface 2 (Figure 2B). Interface 1 allows a classical GST dimerization, where helices α2 and α3 of one monomer interact with α3 and α2 of the second monomer and are parallel one to each other. The alternative interface 1’, was observed in the crystal structure of the N-terminal domain of *Plasmodium vivax* tRip (13), with helices α2 and α3 being oriented perpendicularly to each other’s. As for interface 2, this dimerization involves mainly a stacking interaction between 2 strictly conserved arginines protruding from α7 helices of each monomer. Based on sequence analysis, the formation of interface 2 in *Apicomplexa* GST-like domains is possible only for AIMPs (except that of *Cryptosporidium*), ERSs and the two subunits β and γ of EF1 (Table 1, Figure S2).

### Pre-tests for MSC modeling

We set out to combine our knowledge about dimerization of GST-like domains with deep learning-based protein structure modeling, CollabFold, to identify and determine the structures of *Apicomplexa* MSC assemblies. As a starting point, we chose to predict homodimerization for the *S. cerevisiae* AIMP Arc1p. The crystal structure of Arc1p was determined and is a monomer despite the presence of helices α3 and α7. The five models obtained for this negative control, confirmed that indeed, no homodimer using either interface 1, 1’ or interface 2 could be formed (Figure 3A). We then chose to test the formation of EF1β and γ heterodimers, that is conserved in all eukaryotes and that obviously involve GST-like domains in apicomplexan parasites. As expected, in all models, EF1 β and EF1γ heterodimerize *via* their interface 2 in the absence or in the presence of EF1α and EF1δ within all strains (Figure S3), with pTM scores between 0.41 and 0.74 (Figure S4). Finally, ColabFold was successfully used to model both *Plasmodium* Q- and M- MSCs, and the resulting models were highly consistent with SAXS data and mutagenesis experiments(14,16).

**Figure 3.**
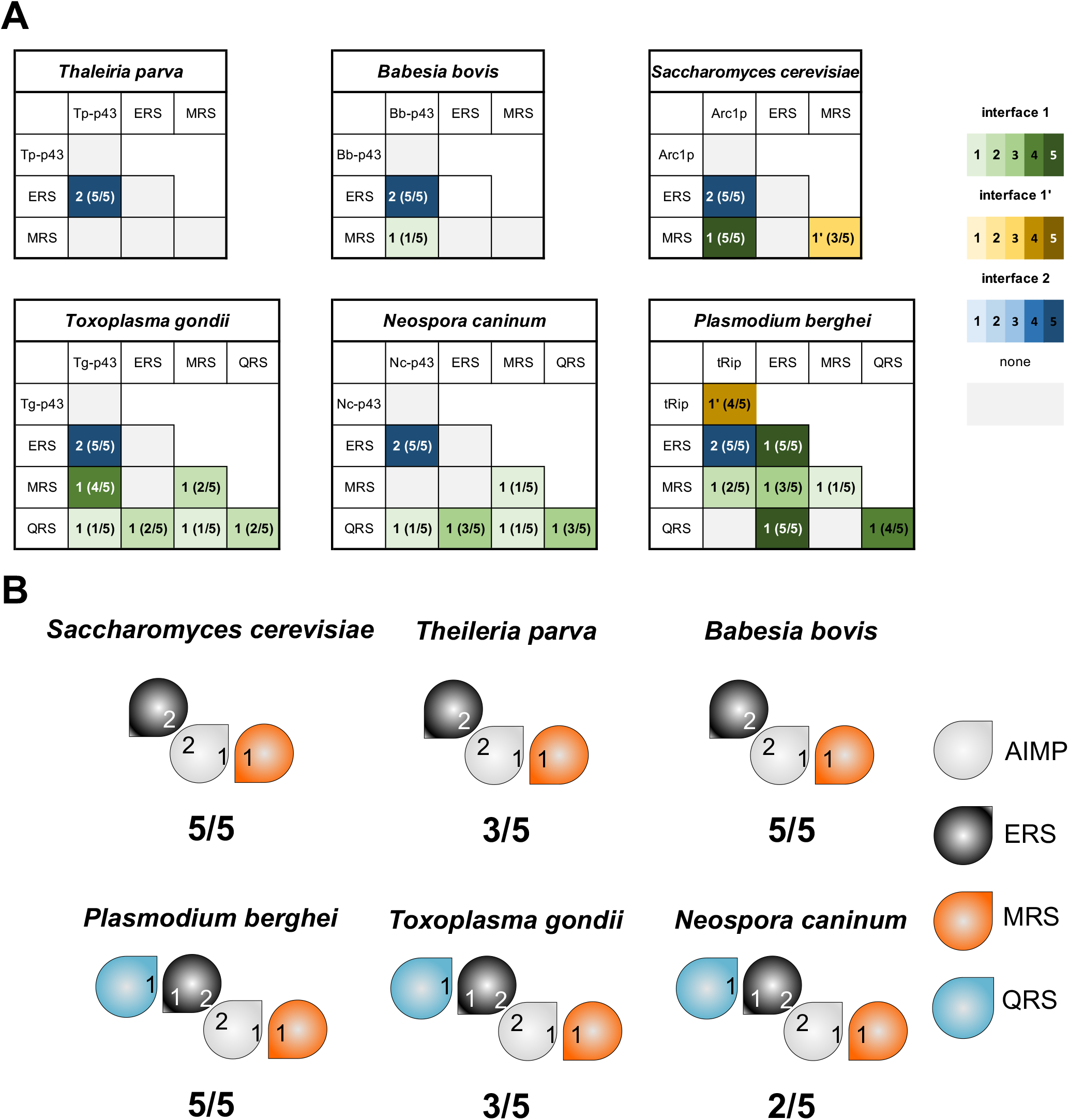
Predicted interactions between MSC GST-like domains in yeast and *Apicomplexa* parasites. (**A**) pairwise interactions. Homo- and hetero-dimers were predicted with ColabFold using the sequences of the GST-like domains involved in yeast and *Apicomplexa* MSCs. For each combination of dimers, the occurrence of canonical GST-like interfaces 1, 1’ or 2 is indicated by a gradient of green, yellow, and blue respectively. The score corresponds to the number of models displaying these interfaces (n = 5). (**B**) ColabFold predictions of heteromeric GST-like complexes are schematized with drop shape, with AIMP colored in grey, ERS in black, QRS in blue and MRS in orange and displaying interfaces 1, 1’ and 2. The *S. cerevisiae, T. parva and B. bovis* MSCs, containing 3 GST-like domains, AIMP binds MRS *via* interface 1 and ERS *via* interface 2. The 4 GST-like domains of *P. berghei, T. gondii* and *N. caninum*, share the same interaction network: AIMP and ERS heterodimerize through interface 2, QRS binds interface 1 of ERS, and MRS interface 1 of AIMP.

Based on these observations, ColabFold was considered suitable for *de novo* modeling of interactions between GST-like domains. We thus used our knowledge about dimerization of GST-like domains with ColabFold modeling to question the structures of other *Apicomplexa* MSCs. For each parasite’s MSC, we modeled (i) all pairwise interactions between the different GST-like domains (Figure 3A) and (ii) the backbone of the MSC in the presence of all GST-like partners (Figure 3B). For each prediction, 5 models were generated, and the relative positions of the domains were explored to identify possible interfaces 1, 1’ and 2. The number of models using only interfaces 1, 1’ and 2 allowed to determine a score (x/5).

### ColabFold predicts that AIMPS et ERS are monomers in other *Apicomplexa* than *Plasmodia*

Since both tRip and ERS homodimerize in *Plasmodium* MSCs *via* interface 1’ or 1, respectively (14). We searched for possible homodimerization of AIMPs’ and ERSs GST-like domains of other Apicomplexa. No model proposed such solutions, suggesting that apart from the GST-like domains of *Plasmodium* tRip and ERS, no other homodimerizes (Figure 3A), which is unexpected for the *Toxoplasma* AIMP, *Tg*-p43, whose homodimerization in solution has been demonstrated (24).

### ColabFold predicts the same interaction network in all *Apicomplexa* MSCs

Based on our sequence analysis, only AIMPs, ERS and MRS are fused to GST-like domains in *Theileria* and *Babesia* genera, a situation that recalls what is known in *S. cerevisiae* where the monomeric AIMP Arc1p binds ERS and MRS (25,26). Consistently, modeling of dimeric interactions (Figure 3A) suggested the formation of the heterodimers AIMP:ERS in both parasites (with a score 5/5). While the formation of the heterodimer AIMP:MRS was possible in *B. bovis* (score 1/5), none was detectable in *T. parva*. Yet, modeling the heterotrimeric GST-like backbone (Figure 3B) allowed the identification of AIMP:MRS subcomplexes in both *T. parva* (3/5) and *B. bovis* (5/5), indicating that both *B. bovis* and *T. parva* MSCs share the same organization than the yeast MSC (Figure S6): AIMP (*Bb*-p43 or *Tp*-p43) binds ERS *via* interface 2 and MRS *via* interface 1. Interestingly, this evolutionary relationship is supported by the fact that *Babesia*, *Theileria* and yeast also possess similar EF1 complexes lacking the sub-unit δ (Table S1, Figure S3).

Van Rooyen and collaborators have shown that in *T. gondii*, 5 proteins associate in a unique cytosolic MSC (24). They correspond to the AIMP *Tg*-p43, ERS, MRS, QRS and the tyrosyl-tRNA synthetase (YRS). Sequence alignments showed that *T. gondii* YRS does not contain a GST-like domain, but rather possesses a N-terminal α helix (1–56), strictly conserved in *N. caninum* (Figure S2G). This helix is absent in *Plasmodia* and has no sequence similarity with other apicomplexan YRSs. Analysis of pairwise interactions clearly indicated heterodimerization of AIMP (*Tg*-p43) and ERS (score 5/5) and of *Tg*-p43 and MRS (4/5) in *T. gondii* (Figure 3A). Prediction of GST-like heterotetramers completed this interaction network: QRS binds ERS *via* interface 1, ERS binds Tg-p43 *via* interface 2 and Tg-p43 binds MRS *via* interface 1 (Figure 3B, Figure S6). Despite being included in modeling, the N-terminal α-helix of YRS showed no convincing interaction with any GST-like domain.

In an unusual way, *T. gondii* and *N. caninum* GST-like sequences are highly conserved, suggesting that they are involved in the formation of similar MSCs. While analysis of pairwise interactions indicated heterodimerization of AIMP (*Nc*-p43) and ERS (5/5), and ERS and QRS (3/5) (Figure 3A), prediction of GST-like heterotetramers revealed that the N. caninum has the same organization than *T. gondii* MSC.

Finally, performing the same tests with the GST-like domains of *Plasmodium*, AIMP (tRip), ERS, QRS and MRS, ColabFold predicted the same interaction network as *Sarcocystidae* MSCs (Figure 3B, Figure S6). However, it has been demonstrated that the formation of a unique MSC containing all 4 proteins is not possible in *Plasmodium* due to the ability of both tRip and ERS to homodimerize with their interfaces 1’ and 1, respectively (Figure 3A). Instead, *Plasmodium* contains two mutually exclusive MSCs, the Q-complex (tRip, ERS and QRS) and the M-complex (tRip, ERS and MRS) (Figure 4A) (14,16). Homodimerization of tRip in the MSC-Q prevents the binding of MRS tRip interface 1 and homodimerization of ERS in the MSC-M prevents the interaction of QRS with ERS interface 1.

**Figure 4.**
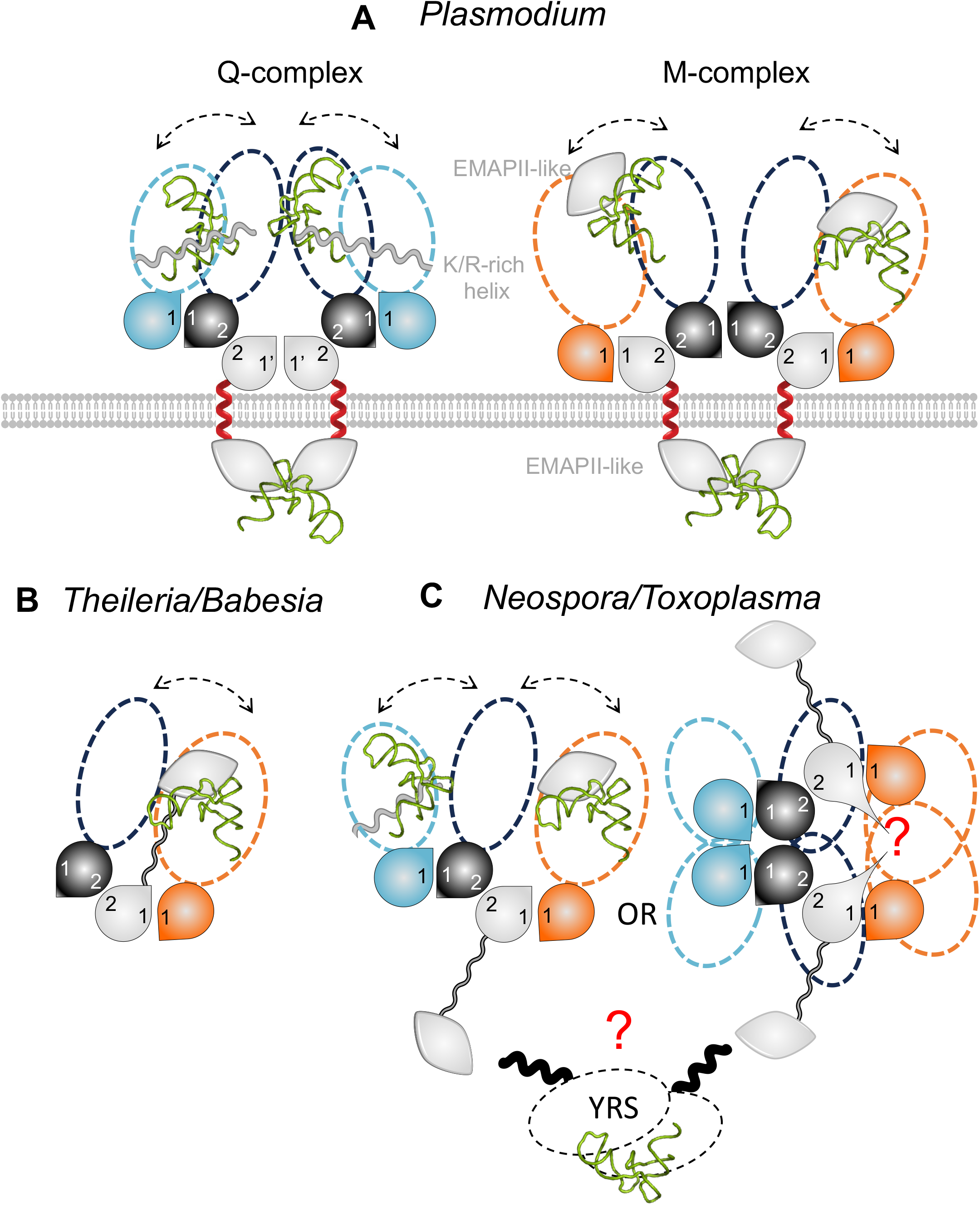
Composition and architecture of *Apicomplexa* MSC complexes. MSCs models of (**A**) *Plasmodia* (14), (**B**) *Theileria* and *Babesia* and (**C**) *Toxoplasma* and *Neospora* are shown. Color code is the same than in the legend of Figure 3. Schematic views of the complexes are built from GST-like domains (drops), minimal core enzymes (catalytic domain and the anticodon-binding domain) and additional RNA binding domains. An EMAPII-like domain is appended to the C-terminus of MRSs (except in *Theileria* and *Babesia* complexes) and a positively charged helix is attached at the C-terminus of QRSs. These domains provide additional non-specific tRNA binding properties to the aaRSs present in MSCs (14). The EMAP-II like domain of AIMPs could be involved in the binding of different tRNAs, either Host tRNAs in *Plasmodium* (**A**) or tRNA^Glu^ or tRNA^Met^ in *Theileria* and *Babesia* (**B**). In *Toxoplasma and Neospora*, 2 MSCs with different stoichiometries are shown, either 1:1:1:1 or 2:2:2:2 (RNA binding domains are not shown). The YRS is a dimer with 2 N-terminal helical domains (black), but no interaction can be proposed.

### The possibility of homodimerization of Tg-p43 and modeling of 2 T. gondii MSCs

To test whether arrangements similar to those identified in the 2 *Plasmodium* MSCs exist in *Sarcocystidae*, we used two copies of Tg-p43, ERS and QRS or MRS to predict *T. gondii* dimeric MSCs. No model proposed *Tg*-p43 or ERS homodimerization using canonical GST-like interface (Figure 3B), which is consistent with the absence of prediction of *Tg*-p43 or ERS canonical homodimers (Figure 3A). However, whether alone or in the models of dimeric MSCs, *Tg*-p43 homodimerizes always through the same non canonical interface involving the insertion between strands β2 and β3 (Figure S2C) and lead to 2 models (Figure S5) consistent with the interaction network proposed in Figure 3B. If biologically relevant, this Tg-p43 homodimerization pattern would explain the dimer of Tg-p43 observed in solution and still lead to the formation of a unique MSC containing all 4 GST-like partners (24) (Figure 4 C, Figure S7).

### Proposed consequences of MSC formation on tRNA binding

The membrane localization of both *Plasmodium* MSCs results in the presence of the tRip tRNA-binding domain (EMAPII-like) outside the parasite and therefore it cannot participate in the aminoacylation reaction(s) catalyzed by the aaRSs present in the complexes. This is different from what was shown in the yeast complex (27,28). However, we proposed that the tRNA binding domains fused to MRS (EMAPII-like domain) and QRS (positively charged helix) replace the tRip EMAPII-like domain to increase aaRSs affinity for their cognate tRNAs (Figure 4A). However, in *Toxoplasma*, immunolocalization experiments demonstrated that the MSC components are cytosolic. The homology of *Theileria* and *Babesia* MSCs with the yeast complex (as well as a very low probability for the presence of a transmembrane helix in the corresponding AIMPs) strongly suggests that these MSCs are also cytosolic. We therefore proposed possible role of each of the tRNA-binding domains associated with the complexes. *T. parva* and *B. bovis* MSCs contain only one tRNA binding domain in the AIMP, a situation that mimics the yeast configuration: The EMAPII-like domain of the AIMP would augment the affinity of ERS and MRS for their cognate tRNAs, thus increasing their aminoacylation efficiency (Figure 4B). As for *T. gondii* and *N. caninum* MSCs, they contain many additional binding domains, on the AIMP and both the MRS and QRS. One can propose that MRS and QRS tRNA binding domains participate to their respective aminoacylation and to glutamylation (Figure 4C) and the AIMP EMAPII-like domain could help tyrosylation.

### What makes *Plasmodium* translation so different that it needs tRNA import *via* membrane associated MSCs?

MSCs predictions indicate that unlike *Plasmodium*, other *Apicomplexa* parasites have only one cytosolic MSC. These observations suggest that *Plasmodium* is thus the only *Apicomplexa* parasite with membrane associated MSCs and which requires a tRip-directed tRNA import for an optimal protein synthesis. Therefore, we inventoried the nuclear tRNA genes for each of the selected species of the *Apicomplexa* phylum (Table 2). Apicomplexan nuclear genomes contain a relatively small number of tRNA genes, compared to other protozoa (http://gtrnadb.ucsc.edu/). When analyzing the *P. falciparum* genome, authors had already noticed that this organism has few tRNA genes, with only one gene copy for a given anticodon (isoacceptor) (29,30). The only exception being tRNA^Met^ (CAT), which is logically present in 2 copies: one copy for translation initiation and one copy for elongation. Here, we observe that these features are well conserved not only in other *Plasmodia* but also in other *Apicomplexa* such as *Babesia*, *Cryptosporidium* and *Theileria* which all have between 45 and 51 tRNA genes. In these parasites, the distribution of isoacceptors is homogeneous with few exceptions: *B. bigemina* which encodes more tRNA isodecoders than the other two *Babesia* species. Moreover, the tRNA^Sec^ is found only in *Plasmodia*, *Neospora*, and *Toxoplasma* and is absent from the genomes of *Babesia*, *Cryptosoporidium*, and *Theileria*; This was confirmed by the joint absence of the genes encoding SelB (selenocysteine-tRNA-specific elongation factor) or SelD (Selenophosphate Synthase), two proteins that belong to the specific tRNA^Sec^ machinery and allow selenocysteine insertion into proteins (31).

**Table 2.**
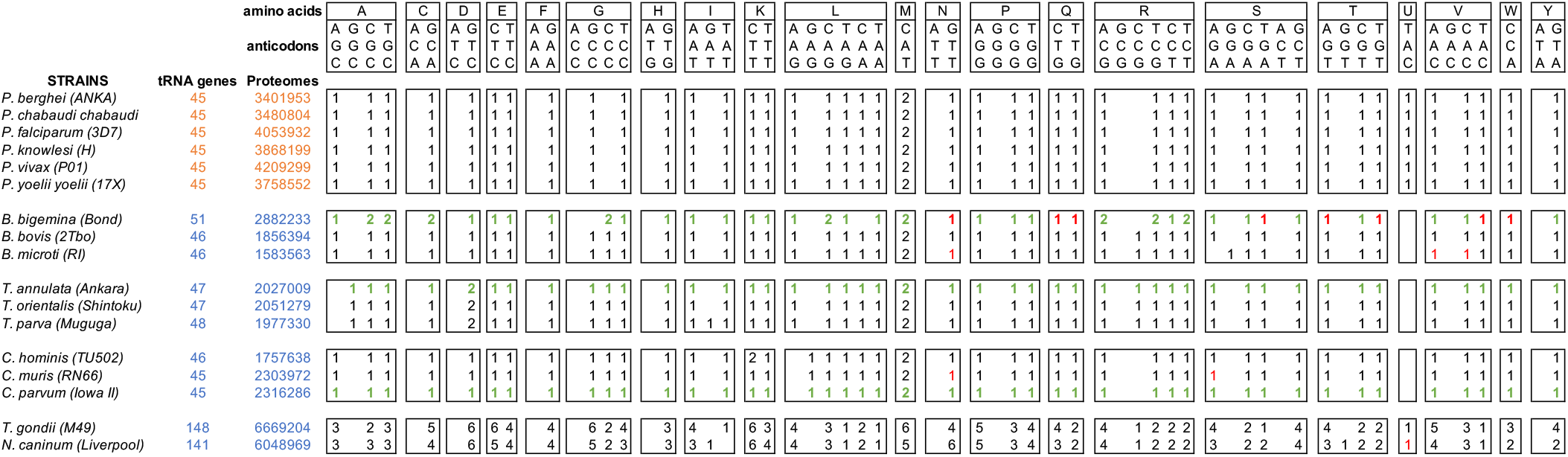
tRNA genes present in the selected *Apicomplexa* genomes. The tRNA genes were retrieved directly from annotated genomes of *Plasmodia*, *B. bovis*, *B. microti*, *T. orientalis*, *T. parva*, *C. hominis*, *C. muris*, *T. gondii* and *N. caninum* (EupathDB). In *B. bigemina*, *T. annulata* and *C. parvum*, tRNA genes were identified using the tRNAscan-SE tRNA gene detection program (green). Some “missing” tRNAs in the genomes of *B. bigemina*, *B. microti*, *C. muris* and *N. caninum* were found manually by Blast with a sequence from an organism of the same genus (in red).

*Plasmodia*, *Cryptosporidium*, *Theileria* and *Babesia* genomes contain roughly one gene copy for each tRNA isoacceptor. This is theoretically enough to support the decoding of their complete proteome. In *Toxoplasma* and *Neospora*, the number of tRNA genes is much higher, with several copies of each tRNA isoacceptor. Despite the difference in tRNA content, there is an excellent correlation (R = 0.97) between the number of nuclear tRNA genes and the size of the corresponding proteomes (total number of encoded amino acids), except for *Plasmodia* (Figure 5A). This suggests that the number of tRNA genes is not adapted to the size of the genome to be translated in *Plasmodia*.

**Figure 5.**
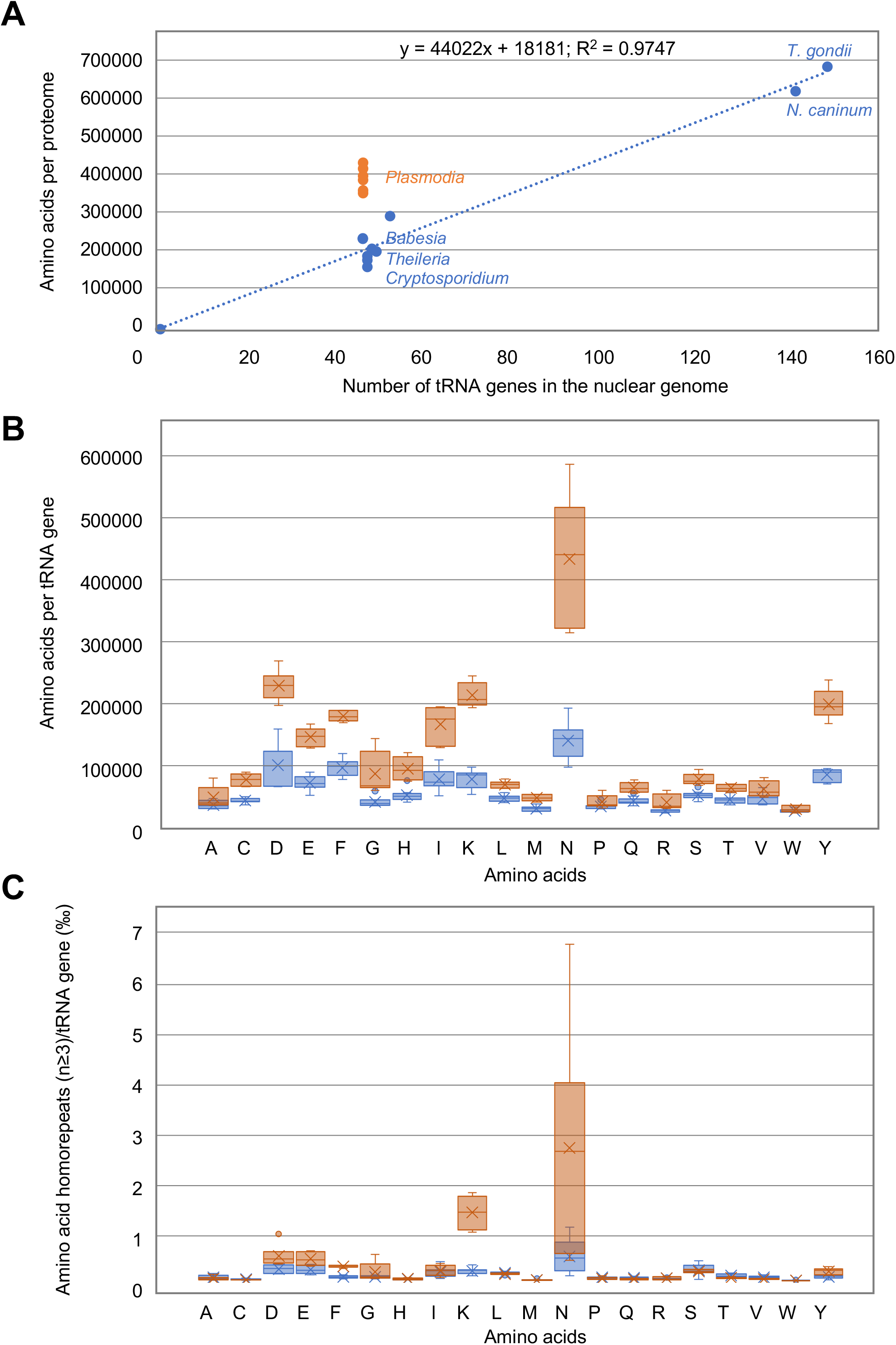
Comparison of tRNA usage in *Apicomplexa* parasites. (**A**) Correlation between proteome size and the number of tRNA genes in *Plasmodia* (orange) and other *Apicoplexa* (Blue). Data are from Table 2. (**B**) tRNA usage displayed as amino acids decoded per tRNA gene (all isoacceptors together) in *Plasmodia* (orange) and other *Apicomplexa* parasites (blue); Some *Plasmodia* tRNAs (Asp, Ile, Lys, Asn and Tyr) are potentially highly utilized during translation, compared to their homologues in other *Apicomplexa* parasites. (**C**) tRNA usage displayed as amino acids homorepeats decoded per tRNA gene (all isoacceptors together).

### Correlation between amino acid usage and tRNA availability in *Apicomplexa* parasites

The sizes of the 6 *Plasmodia* proteomes are homogenous with an average of 3.8+/- 0.3 M amino acids (aa). The other *Apicomplexa* genomes can be classified into two categories: (1) small proteomes corresponding to *Cryptosporidium*, *Babesia* and *Theileria* species with sizes ranging from 1.5 to 2.8 M aa (among the 3 *Babesia* species, *B. bigemina* has a proteome with twice the size of the two others) and (2) large proteomes corresponding to *N. caninum* (6 M aa) and *T. gondii* (6.7 M aa) (Table 2).

To compare the tRNAs most used to synthesize proteins among *Apicomplexa* parasites, the ratio of the usage of each aa to the number of tRNA genes that allow their decoding was calculated. In Figure 5B, the higher the box, the more tRNA is needed to decode the genome. The resulting values represent the frequency of tRNA usage, with the largest values corresponding to the most recruited tRNAs in gene translation for a given amino acid. The frequency of tRNA usage is dramatically different in *Plasmodia* and other *Apicomplexa*. The most significant differences occur in the usage of tRNA^Asp^, ^Lys^, ^Asn^ and ^Tyr^. Especially, there is extreme use of tRNA^Asn^ in *Plasmodia*, which correlates with the presence of long asparagine homorepeats (as well as lysine homoreapeats) specific to *Plasmodia* proteins (Figure 5C). However, while asparagine usage is rather homogenous in *Plasmodia* strains, this is not the case for asparagine homorepeats that are scarcer in *P. knowlesi* and *P. vivax*.

## DISCUSSION

In eukaryotes, aaRSs and AIMPs are moonlighting proteins. They participate to tRNA charging when located into the MSC (32–34) and to diverse non-translational, yet crucial, functions activated by MSC release ((35,36) and reviewed in (37,38)). Among the *Apicomplexa* parasites, only the composition of *T. gondii and P. berghei* MSCs has been experimentally established (16,24). While, *T.gondii* has a single cytosolic MSC made up of one AIMP (*Tg*- p43) and four aaRSs (ERS, QRS, MRS and YRSs) (24), *Plasmodium* harbors two distinct MSCs whose structures have been determined in solution (14,16); Both *Plasmodium* complexes are constituted of a dimer of heterodimer tRip:ERS that associates with either QRS or MRS (Figure 4A). Putative MSCs were identified in all *Apicomplexa* parasite considered in this study, except for *Cryptosporidium*. They all contain one AIMP, ERS and MRS with or without an additional QRS. Such MSCs that contain 3 or 4 GST-like domains were more challenging to model (Figure 4 and Figure S5) than the heterodimer that forms between EF1β and γ subunits (Figure S3). However, amongst MSC models, even those without correct interfaces, all are built on a core consisting of an AIMP:ERS heterodimer that forms *via* the interface 2 on which MRS and/or QRS are attached (Figure 3B, Figure S6).

Several observations support the fact that the *Plasmodium* AIMP tRip, combined to one or both MSCs, is involved in tRNAs import inside the parasite and that their import could compensate for the deficiency of some *Plasmodium* tRNAs (15,17). The deletion of the gene encoding tRip not only leads to a reduction in overall protein synthesis, but also to a selective drop of asparagine-rich proteins in the parasite. Also, the (human) tRNA^Asn^ is one of the favorite binders of (*P. falciparum*) tRip, which increases its chances of import (39). However, based on the present study, this tRNA import seems to be restricted to *Plasmodia* only. Indeed, the lack of resemblance between modeled MSCs and Plasmodium MSCs suggests that they are neither membrane-bound nor involved in tRNA import.

The nuclear genomes of *Plasmodia* encode a limited number of tRNA genes. *Cryptosporidium*, *Babesia* and *Theileria* have also single copies of tRNA genes which corresponds to the small size of their proteomes to be produced, whereas *Toxoplasma* and *Neospora* have multiple copies of specific tRNA isoacceptors correlating with the synthesis of larger proteomes. However, this interplay does not exist in *Plasmodia*, whose tRNA genes are single copies but which have larger proteomes to synthesize (Table 2, Figure 5A). Coding for a single copy for each tRNA isoacceptor differs from what is observed in most eukaryotes, where tRNA genes are present in multiple copies; On the one hand the number of tRNA genes is related to the frequency of codon usage in the genome (40) and on the other hand, the relative abundance of each amino acid in the proteome is not random. These rather intricate balances reflect a subtle interplay between the availability of aminoacylated tRNAs and the regulation of expression of well-folded proteins. The correlation between amino acid usage and tRNA gene content is of particular interest in the context of parasitic organisms. Especially, *Plasmodia* amino acid usage is remarkable compared to other *Apicomplexa* parasites, which may be explained partially by the high AT content of some of their genomes (41), but not exclusively. Figure 5B shows that the correlation between amino acid usage and the number of tRNA genes encoded by *Plasmodia* genomes is less homogeneous than what is observed for other *Apicomplexa* and other organisms in general (40). Indeed, the *Plasmodia* strains included in this study are characterized by more amino acids to be polymerized than the corresponding tRNAs could decode (Figure 5A); It appears that specific tRNAs, especially tRNA^Asn^, could be very limiting for the decoding of *Plasmodia* genomes (Figure 5B), independently of the presence of long asparagine homorepetitions in proteins (Figure 5C) and the AT content of the genome.

It is accepted that changes in tRNA gene content can lead to changes in amino acid utilization, which in turn can lead to changes in protein structure and function (42). In this respect, increased utilization of a particular amino acid should lead to an increased number of copies of the corresponding tRNA gene. Clearly, this is not the case in *Plasmodium* where asparagine usage is disconnected from the number of genes encoding tRNA^Asn^. This imbalance could lead to selective pressure on proteins that use a lot of asparagine in their sequences and be beneficial in certain environments or under certain conditions. By elucidating the mechanisms underlying this relationship, we begin to better understand the evolution of *Plasmodium* genomes and pathogenicity/development. Indeed, what better way to control the production of specialized proteins than through the control of host tRNAs import? Import of specific tRNAs could trigger mechanisms adapted to each stage of the parasite development, depending on the tRNA content of the different host cells (43). We propose that the pool of tRNAs available in the host cells and eventually imported into the parasite is a kind of GPS. Import of host tRNAs would modify the concentrations of the different tRNAs present at a given time inside the parasite. Therefore, depending on the nature of the host tRNAs available and preferentially imported, the translation of one or more key regulators could vary and control the development of the parasite by indicating where it is in its developmental cycle and when to move on to the next stage.

## AUTHORS CONTRIBUTIONS

JRJP and MF managed the conception, design, and interpretation of the data, and wrote, reviewed, and edited the manuscript. MF was responsible of fundings. Both authors approved the final article.

## ACKNOWLEDGEMENTS

We thank Julien Lision for initiating the search for tRNA sequences.

