## Supplemental Table and Figures for "*Plasmodium*, the *Apicomplexa* outlier when it comes to protein synthesis"

### Table S1. Annotation of protein genes in EupathDB

| | AIMPs and AARSs | | | | | Translation factors | | | |
|---|---|---|---|---|---|---|---|---|---|
| | AIMP | ERS | MRS | QRS | YRS | EF1-β | EF1-γ | EF1-α | EF1-δ |
| *P. berghei* | PBANKA_1306200 | PBANKA_1362000 | PBANKA_0518700 | PBANKA_1346600 | PBANKA_1222800 | PBANKA_0814200 | PBANKA_1352000 | PBANKA_1133300 | PBANKA_1218900 |
| *P. chabaudi* | PCHAS_1309400 | PCHAS_1366600 | PCHAS_0518800 | PCHAS_1351200 | PCHAS_1223400 | PCHAS_0814500 | PCHAS_1356600 | PCHAS_1133800 | PCHAS_1219600 |
| *P. falciparum* | PF3D7_1442300 | PF3D7_1349200 | PF3D7_1034900 | PF3D7_1331700 | PF3D7_0807900 | PF3D7_0913200 | PF3D7_1338300 | PF3D7_1357100 | PF3D7_0319600 |
| *P. knowlesi* | PKNH_1240000 | PKNH_1251700 | PKNH_0620100 | PKNH_1268800 | PKNH_0110800 | PKNH_0711100 | PKNH_1263000 | PKNH_1114500 | PKNH_0821600 |
| *P. vivax* | PVP01_1264700 | PVP01_1205600 | PVP01_0620000 | PVP01_1222900 | PVP01_0112500 | PVP01_0711600 | PVP01_1216400 | PVP01_1114900 | PVP01_0820600 |
| *P. yoelii* | PY17X_1310000 | PY17X_1367700 | PY17X_0519800 | PY17X_1351500 | PY17X_1226000 | PY17X_0817500 | PY17X_1357200 | PY17X_1134900 | PY17X_1222100 |
| *B. bigemina* | BBBOND_0102100 | BBBOND_0404120 | BBBOND_0207070 | BBBOND_0307760 | BBBOND_0202210 | BBBOND_0110635 | BBBOND_0200780 | BBBOND_0108320 | |
| *B. bovis* | BBOV_III010240 | BBOV_III011840 | BBOV_I001970 | BBOV_III006400 | BBOV_II002250 | BBOV_IV003370 | BBOV_II000890 | BBOV_IV010620 | |
| *B. microti* | BMR1_01G02705 | BmR1_04g04960 | BMR1_02g02000 | BMR1_03g03040 | BMR1_01G00295 | BMR1_03g01425 | BMR1_02g03070 | BMR1_03g03725 | |
| *T. annulata* | TA11475 | TA17035 | TA05325 | TA12530 | TA08075 | TA16305 | TA08705 | TA06720 | |
| *T. orientalis* | TOT_020000112 | TOT_010001344 | TOT_030000712 | TOT_020000303 | TOT_040000556 | TOT_010001036 | TOT_040000424 | TOT_010000685 | |
| *T. parva* | TpMuguga_02g00124 | TpMuguga_01g01053 | TpMuguga_03g00137 | TpMuguga_02g00322<br>TpMuguga_02g02375 | TpMuguga_04g00367 | TpMuguga_01g01096 | TpMuguga_04g00492 | TpMuguga_01g00726 | |
| *C. hominis* | Chro.40444 | Chro.80098 | Chro.80398 | Chro.10244 | Chro.60344 | Chro.20423 | Chro.70492 | Chro.60459 | |
| *C. muris* | CMU_018940 | CMU_024700 | CMU_006170 | CMU_028890 | CMU_031000 | CMU_003010 | CMU_014590 | ? | |
| *C. parvum* | cgd4_3880 | cgd8_790 | cgd8_3460 | cgd1_2130 | cgd6_2970 | cgd2_3950 | cgd7_4450 | cgd6_3990 | |
| *T. gondii* | TGME49_223140 | TGME49_263870 | TGME49_289300 | TGME49_217460-t26 | TGME49_251880 | TGME49_226410 | TGME49_300140 | TGME49_294800 | TGME49_219140 |
| *N. caninum* | NCLIV_006150 | NCLIV_024090 | NCLIV_041650 | NCLIV_062220 | NCLIV_066920 | NCLIV_046550 | NCLIV_064580 | NCLIV_014180 | NCLIV_061340 |

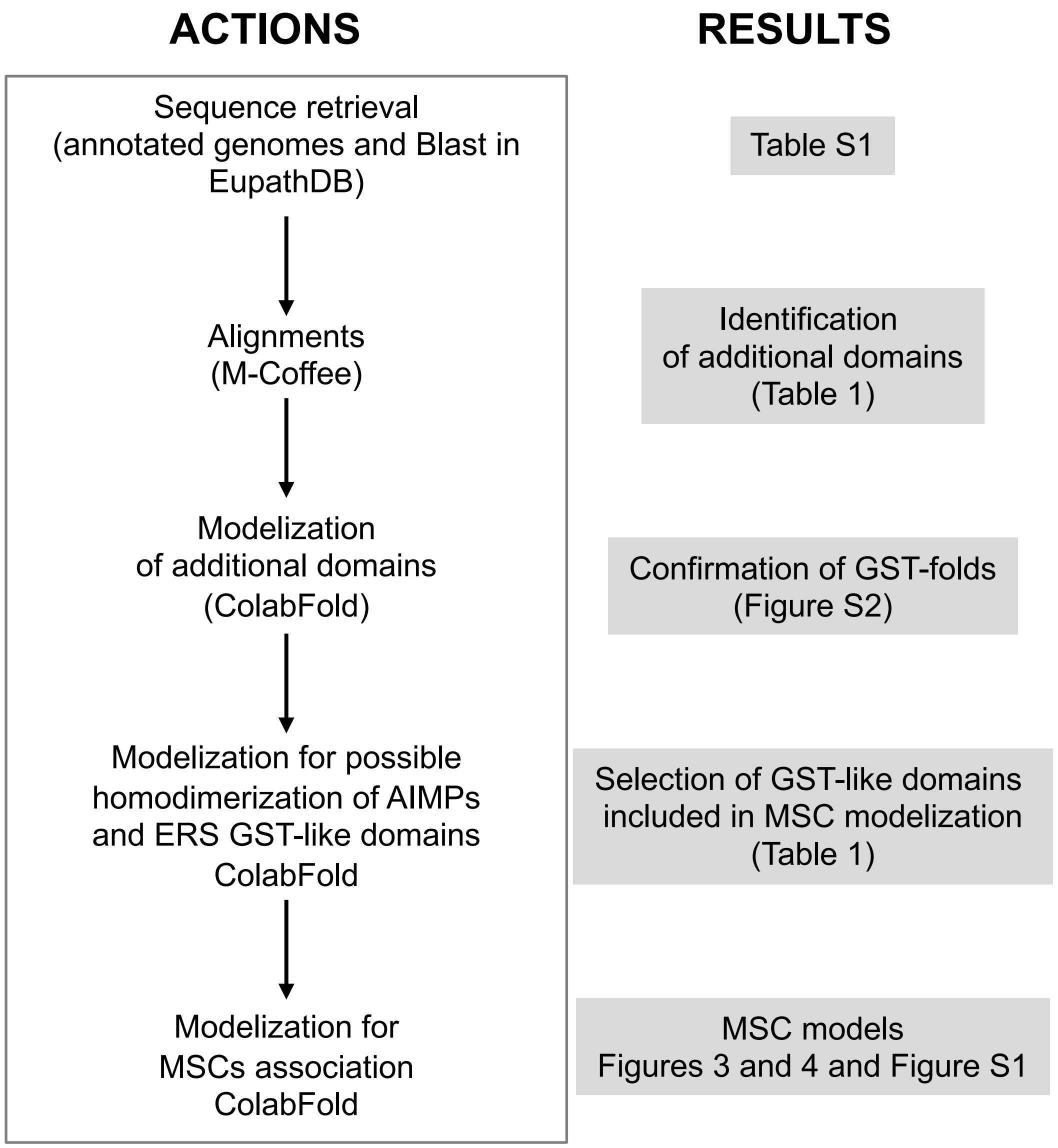


**Figure S1: Pipeline for modeling GST-like interactions in MSCs.** Global approach used to identify GST-like folds in AIMPs and aaRSs and model their association in MSCs with ColabFold. The occurrence of canonical interfaces 1, 1’ and 2 in the resulting models suggest a positive interaction between GST-like domains.

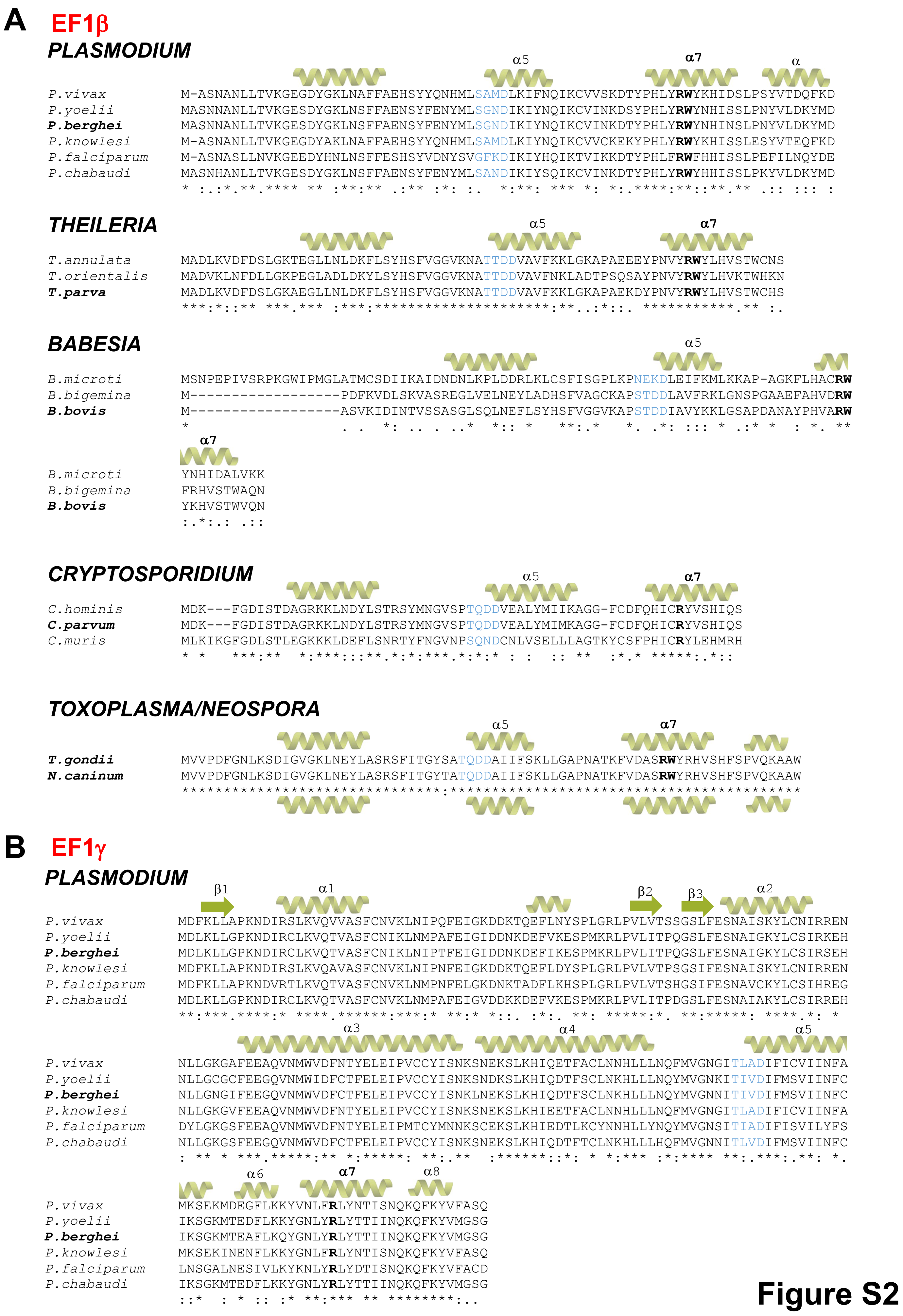

A EF1β
PLASMODIUM
α5 α7 α
P.vivax M-ASNANLLTVKGEGDYGKLNAFFAEHSYYQNHMLSAMDLKIFNQIKCVVSKDTYPHLYRWYKHIDSLPSYVTDQFKD
P.yoelii MASNNANLLTVKGESDYGKLNSFFAENSYFENYMLSGNDIKIYNQIKCVINKDSYPHLYRWYNHISSLPNYVLDKYMD
P.berghei MASNNANLLTVKGESDYGKLNSFFAENSYFENYMLSGNDIKIYNQIKCVINKDTYPHLYRWYNHINSLPNYVLDKYMD
P.knowlesi M-ASNANLLTVKGEGDYAKLNAFFAEHSYYQNHMLSAMDLKIYNQIKCVVCKEKYPHLYRWYKHISSLESYVTEQFKD
P.falciparum M-ASNASLLNVKGEEDYHNLNSFFESHSYVDNYSVGFKDIKIYHQIKTVIKKDTYPHLFRWFHHISSLPEFILNQYDE
P.chabaudi MASNHANLLTVKGESDYGKLNSFFAENSYFENYMLSANDIKIYSQIKCVINKDTYPHLYRWYHHISSLPKYVLDKYMD
THEILERIA
α5 α7
T.annulata MADLKVDFDSLGKTEGLLNLDKFLSYHSFVGGVKNATTDDVAVFKKLGKAPAEEEYPNVYRWYLHVSTWCNS
T.orientalis MADVKLNFDLLGKPEGLADLDKYLSYHSFVGGVKNATTDDVAVFNKLADTPSQSAYPNVYRWYLHVKTWHKN
T.parva MADLKVDFDSLGKAEGLLNLDKFLSYHSFVGGVKNATTDDVAVFKKLGKAPAEKDYPNVYRWYLHVSTWCHS
BABESIA
α5
B.microti MSNPEPIVSRPKGWIPMGLATMCSDIIKAIDNDNLKPLDDRLKLCSFISGPLKPNEKDLEIFKMLKKAP-AGKFLHACRW
B.bigemina M------------------PDFKVDLSKVASREGLVELNEYLADHSFVAGCKAPSTDDLAVFRKLGNSPGAAEFAHVDRW
B.bovis M------------------ASVKIDINTVSSASGLSQLNEFLSYHSFVGGVKAPSTDDIAVYKKLGSAPDANAYPHVARW
α7
B.microti YNHIDALVKK
B.bigemina FRHVSTWAQN
B.bovis YKHVSTWVQN
CRYPTOSPORIDIUM
α5 α7
C.hominis MDK---FGDISTDAGRKKLNDYLSTRSYMNGVSPTQDDVEALYMIIKAGG-FCDFQHICRYVSHIQS
C.parvum MDK---FGDISTDAGRKKLNDYLSTRSYMNGVSPTQDDVEALYMIIKAGG-FCDFQHICRYVSHIQS
C.muris MLKIKGFGDLSTLEGKKKLDEFLSNRTYFNGVNPSQNDCNLVSELLLAGTKYCSFPHICRYLEHMRH
TOXOPLASMA/NEOSPORA
α5 α7
T.gondii MVVPDFGNLKSDIGVGKLNEYLASRSFITGYSATQDDAIIFSKLLGAPNATKFVDASRWYRHVSHFSPVQKAAW
N.caninum MVVPDFGNLKSDIGVGKLNEYLASRSFITGYTATQDDAIIFSKLLGAPNATKFVDASRWYRHVSHFSPVQKAAW
B EF1γ
PLASMODIUM
β1 α1 β2 β3 α2
P.vivax MDFKLLAPKNDIRSLKVQVVASFCNVKLNIPQFEIGKDDKTQEFLNYSPLGRLPVLVTSSGSLFESNAISKYLCNIRREN
P.yoelii MDLKLLGPKNDIRCLKVQTVASFCNIKLNMPAFEIGIDDNKDEFVKESPMKRLPVLITPQGSLFESNAIGKYLCSIRKEH
P.berghei MDLKLLGPKNDIRCLKVQTVASFCNIKLNIPTFEIGIDDNKDEFIKESPMKRLPVLITPQGSLFESNAIGKYLCSIRSEH
P.knowlesi MDFKLLAPKNDIRSLKVQAVASFCNVKLNIPNFEIGKDDKTQEFLDYSPLGRLPVLVTPSGSIFESNAISKYLCNIRREN
P.falciparum MDFKLLAPKNDVRTLKVQTVASFCNVKLNMPNFELGKDNKTADFLKHSPLGRLPVLVTSHGSIFESNAVCKYLCSIHREG
P.chabaudi MDLKLLGPKNDIRCLKVQTVASFCNVKLNMPAFEIGVDDKKDEFVKESPMKRLPVLITPDGSLFESNAIAKYLCSIRREH
α3 α4 α5
P.vivax NLLGKGAFEEAQVNMWVDFNTYELEIPVCCYISNKSNEKSLKHIQETFACLNNHLLLNQFMVGNGITLADIFICVIINFA
P.yoelii NLLGCGCFEEGQVNMWIDFCTFELEIPVCCYISNKSNDKSLKHIQDTFSCLNKHLLLNQYMVGNKITIVDIFMSVIINFC
P.berghei NLLGNGIFEEGQVNMWVDFCTFELEIPVCCYISNKLNEKSLKHIQDTFSCLNKHLLLNQYMVGNNITIVDIFMSVIINFC
P.knowlesi NLLGKGVFEEAQVNMWVDFNTYELEIPVCCYISNKSNEKSLKHIEETFACLNNHLLLNQFMVGNGITLADIFICVIINFA
P.falciparum DYLGKGSFEEAQVNMWVDFNTYELEIPMTCYMNNKSCEKSLKHIEDTLKCYNNHLLYNQYMVGNSITIADIFISVILYFS
P.chabaudi NLLGKGSFEEGQVNMWVDFCTFELEIPVCCYISNKSNEKSLKHIQDTFTCLNKHLLLHQFMVGNNITLVDIFMSVIINFC
α6 α7 α8
P.vivax MKSEKMDEGFLKKYVNLFRLYNTISNQKQFKYVFASQ
P.yoelii IKSGKMTEDFLKKYGNLYRLYTTIINQKQFKYVMGSG
P.berghei IKSGKMTEAFLKQYGNLYRLYTTIINQKQFKYVMGSG
P.knowlesi MKSEKINENFLKKYGNLFRLYNTISNQKQFKYVFASQ
P.falciparum LNSGALNESIVLKYKNLYRLYDTISNQKQFKYVFACD
P.chabaudi IKSGKMTEDFLKKYGNLYRLYTTIINQKQFKYVMGSG

Figure S2

#### THEILERIA

```
                β1          α1            β2                     β3      β4      α2
T.annulata      MVRKVVGVDLEDYGTKSVLVTAAWANVNFDFEVAKGTCCASQAKKNAFSELKHAVTLVLDKDLGTFCGHVTVCRHLLDHV
T.orientalis    --MKVVGVDVDDYGTKCVLVTAAWSDLKFDFEASKGVCCSSTCEKDAFSELKHAVTLVQDNDLGTFCGHVSVCRHLMDLV
T.parva         --MKVVGVDVDDYGTKSVLVTAAWANLNFDFEVAKGTCCASQAKKNAFSELKHAVTLVLDKDLGTFCGHVTVVRHLLDQV
                  ******::*****.*******::::****.:**.**:*  .:*:************ *:*********:* ***:* *

                           α3                                 α4                  α5
T.annulata      SKGTDEKSAFERSDTSSWVEFFYSRLDFETKLDKSSVVTRGSNSDLSKVVVSLNNYLATKTFLVAEKLSPADLVCAVSLD
T.orientalis    CRGTDDKAVFVKGDTSSWLEFFYLRLEMESKLDTKSVVTKGDNSDLSKVVHAMNKYLATATFLVCEKLGPADLVFAVTLE
T.parva         SKGTDEKSAFERSDTSSWVEFFYSRLDFETKLDKSSVVTKGSNSDLSKVVNSLNNYLATKTFLVAEKLSPADLVCAVSLD
                .:***:*:.* :.*****:**** **::*:***..****:*.******** ::*:**** ****.***.***** **:*:

                       α6   α7          α8
T.annulata      HLVHQKRLDAEYLEKELLHLARFLRTVKASDSFQKFLKMVENYKST
T.orientalis    HLMHHKRLDAAYLEAELLHVARFLRTVKASDKYQKFLEMVERFRGA
T.parva         HLVHQKRLDPEYLEKELLHLARFLRTVKASEAYQKFLKMVENYKSN
                **:*:****. *** ****:**********: :****:***.::.
```

#### BABESIA

```
                                  β1         α1        β2                  β3
B.microti       -----------------MQIIGD-NNSVQTKIAVALTVFSGH--DPKVLIGNGNETSLLPQSF-KLMPMLKEST-----N
B.bigemina      MVGCASAAAALTSLTHLQKVIGVDFNDLGTKNVLAAAQFLGVAYDFQLSKGSCCSSEGKKDEFSEVKPCVSVELCGGLGA
B.bovis         -----------------MKIIGVDLTDLGTKNVLAAAAFYNVSYDFTLAKGGCCSSEGKKDEFSDVKPSVSVELSGGLGS
                                  ::**    ..: ** .:* : * .    *  :  *.  .:.   :.* .: * :. .

                β4     α2                   α3                                          α4
B.microti       HVGTLTIYNYLSGKYAKNFFGATTETRNQITTLFDY------------SVTNLMKYLHLPLGDCLKNKLSELNKCFLKNT
B.bigemina      FCGSISICRHMMEQKKKGDSPAELFARSDATSWMEFFYDRFDVGSRCQAVTQVDASSTSELAKCLR----ATDAFLLSRT
B.bovis         FCGSISISRHLMEEHKKGSAPEELFARSDITSWMEFFYERLDVGSRCQAVTKVVHGDHSELAKCIR----AIDAFLLTHT
                . *:::* .::   :   *.        :*.: *: :::             :**::        *..*::       :  :*..*

                              α5               α6       α7            α8
B.microti       FFVEDQITAADVSMAIIIEELLSRNIIDNCIMENDYYCVKRWFETVTAQDLYKKY---LCTLKGASS
B.bigemina      FLVGESLSVADIVCAVTVKHLVSIGKLEAGYLRTGLRNLSRLIRTVEASDAFVNYEKKLASLQSGSA
B.bovis         FLVGEALSLADVVCAITVDHLISISKLDTEYLASRLLNLSRLIRTVKESDVYAAYHKKLATLNSGVA
                *:* : :: **:  *: :..*:* . ::   :  .    :.* :.**  .* :  *   *.:*:..
```

#### CRYPTOSPORIDIUM

```
                   β1         α1        β2                 β3     β4      α2
C.hominis       --MVYTLFGLDSCPLVSIIKSVAEITGDKISHKVSDGTDHKDLDFHLTF-GTPVLDTQFGKIFGTSTILRFLARGSPYTN
C.parvum        --MVYTLFGLDSCPLVSIIKSVAEITGDKISHKVSDGTDHKDLDFHLAF-GTPVLDTQFGKIFGTSTILRFLARGSPYTN
C.muris         MNTEHKVYYVMKKCPLSVVIASVAEIAQVKVKVIDRNEFTEERQSSLIFGDKTPVISTPFGDIFGIHSILRYLARCAPASF
                    :.:: :..***  :* *****:  *:.     :    .:   .   :     ***:.* **.***  :***:***  :* :

                            α3                                α4                    α5
C.hominis       LYGRTDSDATHVDMILQYIDTTLSSLVVPLFKKSGD--VKSYVSE--LTKIFESFNKYFNYRTFVAGERITIADILLATI
C.parvum        LYGRTDSDATHVDMILQYIDTTLSSLVVPLFKKSGD--VKSYGSE--LTKIFESFNKYFNYRTFVAGERITIADVLLATI
C.muris         LYGETLYESALVDALLEGFHSNILKPIKALCSTACPSECKSDKSEVDLGTVLSNLDDYLLSRTFLAGHRVTIADILIAVV
                ***.*   ::: ** :*: :.:.: . : .* ..:      **   **   *  .::..::.*:  ***:**.*:****:*:*.:

                          α7          α8
C.hominis       IDYSIKCSAGALSLKSCVNLQRYVDTILAQPSIKKHYGEYVKKCSASKDSQAK
C.parvum        IDCSIKCSAGALSLKSCVNLQRYIDTILAQPSIKKHYGEYVKKCSASKDSQAK
C.muris         ICFTVDISKGSLSINGLVNLSRFVNTIMNNYIIRKHYD-NLKKYQTCTDSNTN
                *   ::. * *:**::. ***.*:::**: :  *:***.  :** .:..**:::
```

#### TOXOPLASMA/NEOSPORA

```
                β1        α1         β2                      β3     β4      α2
T.gondii        MKLLTPKDDVRGRKVQLVAAFLDLPLQTVPFTVGKDDKDPAFLAKSPLGRLPLLESEVGGVCLFESNAICRFLARLRADK
N.caninum       MKLLTPKDDVRGRKVQLVAAFLDVPLQTVPFTVGKDDKDPAFLAKSPLGRLPLLESEVGGVCLFESNAICRFLARLRADK
                **********************:*********************************************************

                             α3                              α4                      α5
T.gondii        CLYGETLAEQGQVDMWLDFSTLEVEIPMCCLVQGGKVAERAQSDLAQALNAVDAHLKTRTFMVGENITIADLCLVAVLSY
N.caninum       CLYGETLAEQGEVDMWLDFSTLEVEIPMCCLVQGGKVAERAQSDLSQALNAVNAHLKTRTFMVGENITIADLCLVTVLSY
                ***********:*********************************:******:**************************:****

                         α6    α7         α8
T.gondii        GFRSGKVDAAALLEKRPYLKRFYETVVNQKSFKKIFGE
N.caninum       GFRSGKVDCAAILGQRPFLKRYYETLVNQKSFKKVFGE
                ********.**:* :**:***:***:*********:***
```

### Figure S2

### C AIMPs

#### *PLASMODIUM* (tRip)

```
              β1    α1                 β2     β3   β4   α2
P.vivax       MCVLTLVQDDVKSDILKLVLDFIKAVVVKDDEKVAFPEVRHEKKISFQYKDKQYKELFCTLYAIIDIYDCYNELFNEDEG
P.yoelii      MCTLNLVEDDIKSDILKLALDFIKTNIVENDDNVVFPEIKYGQKISYEHNNKTYKEFFCSLYAIIDTYNCYSQFFCEDED
P.berghei     MCTLNLVENDIKSDILKLALDFIKTNIVENDDNVVFPEIKYGQKISYEHNKKTYKEFFCSLYAIIDTYNCYSQFFCENED
P.knowlesi    MCVLTLVQDDVKSDILKLVLDFIKAVVIKDNEKVAFPEVRHEKKISFEYKEKQYKELFCTLYAIIDIYNCYNELFNEDEG
P.falciparum  MCVLTLVKDDIKSDILKLVLDYIKVTVVQDNENVKLPEICYDKKITLQYKNKTYKDLFCTLYALIDIYDCYSELFNEDEG
P.chabaudi    MSTLYLVEDDIKSDILNLVLDFIKINIVENDDSVAFPEIKYSQTISYEHNNKTYKEFFCSLYAIIDTYNCYSQFFCEDEN
              *..* **::*:*****:*.**:**  :::::::.* :**: : :.*: :::.* **::**:***:** *:**.::* *:*.

                α3               α4                    α5              α6     α7
P.vivax       KVSENEEFIFHLASDKFKLKQLDMKHLNDLLCEKSYIVSNRHASIVDIFYFCSVYKPLSEMPAKERVEISHIYRWFLHIQ
P.yoelii      KVSESEEFIFNLASDKFKLKPLDMKHLNDILRERSYIVSDKHASIVDIFYFCCVYKILKPMTAKERVEYYHICRWYIHLQ
P.berghei     KVSESEEFIFNLASDKFKLKPLDMKHLNDILRERSYIVSDKHASLVDIFYFCCVYKILKPMTAKEKVEYYHICRWYIHLQ
P.knowlesi    KVSENEEFIFHLASDKFTLKQLDMKHLNDLLCEKSYIVSNRHASIVDIFYFCSVYKLLSEMPPKERVEFSHIYRWFLHIQ
P.falciparum  KVSENEEFIFHLASDKYILKQSDMKHLNDLLCEKSYIISNKHASIVDIFYFCAIHKLLDEMAVKERIEFSYIYRWYLHIQ
P.chabaudi    KVSESEEFIFNLASDKFKLKPLDMKHLNDILRERSYIVSDKHASIVDIFYFCCVYKILKPMAAKEKVEYYHICRWYIHLQ
              ****.*****:*****: **  *******:* *:***:*::***:*******.::* *. *. **::*  :* **::*:*

                        α8
P.vivax       ETLVGKFTTLKKLEVRDSLETFLNSKNV---------------MSPGERANSVSMRQQKDDKNAKNENGDNEKGKKKNNA
P.yoelii      ETLICDFVKLHKLDIRSGVESLLNNRTV---------------TCTDEKVSNEQVSPKKDKPNKKNADSK--DNN-GKNK
P.berghei     ETLICDFVKLHKLDIRSGVESLLNSGIV---------------TCTNEKASNEQVGSKIDKANKKNSDSK--DNN-GKNK
P.knowlesi    ETLVGNFTTLKKLEVRDSLETFLNSKNVVNPSERANNAGAGQQKDDKNVKNANGENAENGQNGQNVQNVQNEKSKKKNNA
P.falciparum  ETLLANFSTLKKLIVKDSLENLLNNKTT----------------NNAPEHKNNFVSKESKENKSQNNESPK--NK-KKDV
P.chabaudi    ETLMCEFVKLHKLDIQSGVESLLNSRII----------------TSTNEKGNNEQMGSKKDKGNKKNADSK-DNS-GKNK
              ***: .* .*:** ::..:*.:**.

P.vivax       QNKNAQ-KKKVEEPKNLDDITRLNIIVGYVEEVEIHPDADTLYCLKINVGEEKSRDICSGLRLKKNSEDLLHKYVLVLAN
P.yoelii      KKNNNAENKDAEESRSLDDISRLNIVVGYVESVEIHSGADTLYCLKVNVGEDQLRDICSGLRNKKNPEDLLNKYVLVLAN
P.berghei     KKSNNAENKDAEDSRSLDDISRLNIVVGYVESVEIHSGADTLYCLKVNVGENQLRDICSGLRNKKNPEDLLNKYVLVLAN
P.knowlesi    QNKNNQ-KKKVEEPKSLDDITRLNIIVGYVEEVEIHPDADTLYCLKINVGEEKSRDICSGLRLKKNSEDLLHKYVLVLAN
P.falciparum  QNKNNAPNKKVEETKKLDDISRLNVLVGYVEQVEIHPDADTLYCLKINLGEDKPRDICSGLRNKKNAEDLLNKYVLVLAN
P.chabaudi    KKNNNAENKDAEETRSLDDISRLNIVVGYVESVEIHSGADTLYCLKVNVGEDQVRDICSGLRNKKNPEDLLNKYVLVLAN
              ::.*   :*..*:.:.****:***::*****.****..********:*:**:: ******** ***.****:********

P.vivax       LKEKSLRGRKSHGMVLCGSFGEQIELLAPPDGVNVGERIICENMDVNKLPDKTLSFDKEKNPFFHIQPHLLVKNGVAHYK
P.yoelii      LKEKSLRGRKSFGMILCGSFEERVELLTPPPGVKIGERITFENMNTNSLPDKTLSSVKEKNAFFLIQPNLLINNGVAFYK
P.berghei     LKEKSLRGRKSFGMILCGSFEERVELLTPPPGVKIGERITFENMNTNSLPDKTLSSVKEKNAFFLIQPNLVINNGVAFYK
P.knowlesi    LKEKSLRGRKSHGMVLCASFGEQIELLAPPSGVKVGERIICENMDVNKLPDKTLSSDKDKNPFFHIQPHLLVKNGVAYYK
P.falciparum  LKEKSLRGKKSHGMVLCGSFDEKVELLVPPNGVKIGERILFHNMDPNVIPDKNLSSDKEKNPFFHIQPHLILKDGVAHYK
P.chabaudi    LKEKLLRGRKSFGMVLCGSFEERVELLVPPPGVKVGERITFENVNTTGLPDKTLSSVKEKNAFFLIQPNFVINNGVAFYK
              **** ***:**.**:**.** *::***.** **::****  .*:: . :***.**  *:**.** ***::::::***.**

P.vivax       DAKWLSSKGEITCPLEQGTIS
P.yoelii      NNKWLSSQGEITCAIEQGTIS
P.berghei     NNKWLSSQGEITCMIEHGTIS
P.knowlesi    DTKWLSSKGEIICPLEEGTIS
P.falciparum  DTKWISSQGDITCVLNQGTIS
P.chabaudi    ENKWLSSQGEITCALDQGTIS
              : **:**:*:* * ::.****
```

#### *THEILERIA* (*Tp*-$p_{42}$)

```
               β1       α1           β2          β3                 α2
T.annulata     MLELTVNSNSQESQISKLLWKYLNLNHATLLFTTDKDTESALKFKDGRETVKHFPLILELVLTTSDTGSLLIPNEPELKG
T.orientalis   MIELVVYSDSQESQIVKLLLKYLNLNDVSLTLSTNKDEELFLKLPGETEPVKHFPLMLERLLKSSPHGSLLLPTDPELKA
T.parva        MLELTVNTNSQESQISKLLWKYLNLNHATLLFTTDNDTESALKFKDGRETVKHFPLILELVLTTSDTGSLLIPNEPELKG
               *:**.* ::****** *** ******..:* ::*::* *  **: .  *.******:** :*.:*  ****:*.:****.

               α3           α4                    α5            α6      α7
T.annulata     TMDSFVDFAFKHNFDVLDVESIRMLDTYLLNETFFAGPTLTLADVVLFVSLLNWTVRSKPKERSEFPNLMRWFDHLQHLP
T.orientalis   TMDSFVDFGFKHGFNVLDVDSLRMLDNYLLNETFFSGPTISLADVVLFVSVTYWTSRSKSKERMEVPNLMRWFDHLQHLP
T.parva        TMDSFVDFAFKHNFDVLDVESIRMLDTYLLNETFFAGPSLTLADVVLFVSVLNWTVRSKPKERSEFPNLMRWFDHLQHLP
               ********.***.*:****:*:****.********:**:::*********:   ** ***.*** *.**************

T.annulata     ELATCFEEFSVINLFDVCGIKLDFI-FRTNSLYRSLRRKKLKRKKKRKK--------NRSFEG----------------
T.orientalis   EFASCFEEFKVIQLFDEKLVMTEPKKGKPRAAYHPSEKAEKKKEKKEKPKVEKPPVETRPYDDVTRLNVVVGLVKSIRKH
T.parva        ELAACFEEFSVINLFDNKLVISQPK--AEKVE--------KKKEKKEKPKIQAPPAETRPYDDVTRLNVVVGSVLSIRKH
               *:*:*****.**:***    :   :    .

T.annulata     ---CRWQCIVNQEAPGCRQSICSGLVDFLAPDDILNKKVCVLANLPKKNLRGEESNGMVLCVSNEDKTSVELLEPPTETP
T.orientalis   EDADKLYCLKID-VGSEVRSICSGLVDFLMPDQILDKKVCVLANLPKKNLRGEESNGMVLCVSNSDKSSVELLEPPVDAP
T.parva        QDADKLYCLKVD-IGTEVRSICSGLVDFLGPDDILNKKVCILANLPKKNLRGEESNGMVLCVSNEGKSSVELLEPPEETP
                   :  *:  :      :********** **:**:****:***********************..*:******** ::*

T.annulata     VGERVNWEGLSGEPDEQLSLKKGKDTFSLVQKDLLCKDNIGYYKVDLFFFKKSF-------SHRNS
T.orientalis   VGERVYWEGYSGEADEQLSSKKGKDTFAMVQKDLNCKENVGFYKVSGA----------------E
T.parva        VGERVSWEGLSGEPDEQLSLKKGKDTFAMVQKDLVCKDNVGYYKSSKFMTSKGPCFCKKISTGTIS
               ***** *** ***.***** *******::***** **:*:*:*:**  .                .
```

**Figure S2**

#### ***BABESIA*** (*Bb*-p43)

```
                 β1       α1             β2        β3   β4   α2
B.microti    MIPLEVHKASNYANIVKLLLSYSNIKDDLIKLTLSDNPQLTLTLPNGTQYTEL-PTIMIYIMGQNDTGKILYPDDNLLK-
B.bigemina   MLEVVVYKDCVEGRLATLLWGYGKLTSA--TLSFAPSRTGMVVKEDGKETPMIWPTLLELAVKDAPTESILVPATEAQKV
B.bovis      MLEIVVYSDCVEGRLARLLWGYGKFNNA--TLSLSSTRTETLVKENGEQSLKQWPDIIEIAVKGASTEDVLVPPTPDQK-
             *: : *:. .  ..:. ** .*.::..    *::: .    :.  :* :     * ::   :    * .:* *     *

                         α3                     α4                  α5            α6
B.microti    ------------AKIDQWISLANKKDYSICDSESIREIDSSLVHTTFLVCNNITLADIVIYSSLLHWAKKSTTKEKELMC
B.bigemina   SALRPVFNFACQRTMYSWIDFAYQRDFSIMESASLKILNDYILNHTFLAGPTVTLADLVVYVSTHSWMMKSEPQDRAEYV
B.bovis      ------------NKMYSWIDFAFQRNFMINQKDSLKILNVYLKPNTFLVGNNVTLADLVAYVSIHSWMLMSEPHDRIEFS
                         .: .**.:* :::: * :. *:: ::  :   ***.  .:****:* * *    *    *  .:::

                 α7                         α8
B.microti    NLSRWYNHIQHLPGISIGNYIDIP---TGTTVSSLSSMLNNVNVCGDN-KKSNKSAQKAEGKYNKKIQNSNFSNDARPVD
B.bigemina   NVVRWFDHIQHLPGIV-NTFKELPLVSVDKDMDVVTAVMEKATVSAAAPAPAGKQAKGQKGAASKKPKDPKPPADDRPIA
B.bovis      NIVRWFDHIQHLPGIV-NTFKDLPLVVVDKDMDLVTAVMDNVTVTAPS---AKNPTKGAIDAKSKKEKSPKPVVDERPIA
             *: **::********    ..: ::*    ...  :.  ::::::..*  .         : : ::   .  .** :..:  * **:

B.microti    DITRLAVRVGLVKEISKHPDADKIYCLKIDIAESKLRDICSGLVEHLKPEEIINKKVCVLSNLKPRKIRGVDSNGMVLCV
B.bigemina   DVSRLNIKVGQIMSVERHPEADKLYCLKVDLGEPELRDICSGLVGYLEPEQIMSQYVCVLSNMKPKSLRGKVSNGMILCV
B.bovis      DVSRLNIRVGQVMSVDRHPEADRLYCLKIDLGEPELRDICSGLVGYLQPDEIMSKYVCVLSNMKPKNLRGRVSNGMVLCV
             *::** ::** : .:.:**:**::****:*:.*.:********* :*:*::*:.: ******:**:.:**  ****:***

B.microti    SYNDK--VELLEPPSDSMVGELITWGDLKGEPDVVLSGKSGKNPFEAVQKDLVCKNKIGYYKDLPFSTLSGVCSCNTLIK
B.bigemina   SGPGKTELELLHPAEGAPVGERIIIEDCVGEPDAVLSTKTGKDPFVAVQPEFNCKDTVAYYKDHRFMTSCGPCRCNSLKA
B.bovis      SNADHTQIELLHPADGTPIGERVTIEGFDGEPDAVLSTKTGKDPFVAVQPDFNCKDTIAYYKDHRFMTTLGPCHCNSFTS
             *   .:   :***.*...: :** :    .    ****.*** *:**:** *** :: **:.:.****  * *   * * **::

B.microti    GTIS
B.bigemina   GTIS
B.bovis      GTIS
             ****
```

#### ***CRYPTOSPORIDIUM***

```
                 β1        α1           β2                   β3                  α2
C.hominis    MAILVKYSKTSNIGKVVSLLVNYLLKQNTELKIEESIIDENSTNEKNINQVVLIKDKENCNDQLENIVDILIYLGSFGKN
C.parvum     MVILVKYSKTSNIGKVVSLLVNYLLKQNTELKIEESIIDENSTNEKNINQVVLIKDKENCNDQLESIVDILIYLGSFGKN
C.muris      MTIKLIINPETSFGTLLILVSNYIINQ----KI--SIGTEYS------NQIASTLILESEGIKLINSVDILKHLGSIYPK
             *.* :   .  :.:*.:: *: **:::*                *       **:.     *.  .  :* . **** :***:  :

                         α3                α4                   α5
C.hominis    LLGKDSSLINESKEIIEKLIKINFNFSKGTEMSEFNEMIKGKSFFLGRSLTIVDLVIYISLYSYFESLKDQNQEYIPEYP
C.parvum     LLGKDSSLINESKEIIEKLIKINFNFSKGTEMSEFNEMIKGKSFFLGRSLTIVDLVIYISLYSYFESLKDQNQDYIPEYP
C.muris      IFKQDMEPVMT-K-FLELLSRDSFNTESDDILQHLEDQISGKSFFFGRFLSLLDLVVYSSIHSYFKSMNNMDSTYTPKYP
             :: :* . :    *  ::* * : .** ... :..::: *.*****:** *:::***:* *::***:*::: :. * *:**

                 α7                    α8
C.hominis    NLSTFFEQIQIVTLIRQSCPEGINYLDLPFFKKKLVKKEKKNTGEHGNNSNHNKQ-ERPLDDPTRIALRVGRILSVERHP
C.parvum     NLSTFFEQIQIVTLIRQSCPEGINYLDLPFFKKKLVKKEKKNTGEHGNNSNNNKQEERPLDDPTRIALRVGRILSVEKHP
C.muris      SLSKFFEFVQIMTPIRQACPENLEYLDLPFFTKKLSNKTKK-----LDNT----SDERSLDDPSRIELRVGKILSVERHP
             .**.*** :**:* ***:***.::*******.*** :* **       :*:     . **.****:** ****:*****:**

C.hominis    TADKLYLEKIDVGEDEPRTILSGLVGVYDLTQKINELVVIVSNLKPRAMRGITSNGMLLCASSAPSGDN--NNSQNQSSK
C.parvum     TADKLYLEKIDVGEDEPRTILSGLVGVYDLTQKINKLVVIVSNLKPRAMRGITSNGMLLCASSAPSGDNNNNNNSQNQSSK
C.muris      SADKLYVEKIDIGEEEPRTILSGLVGIYDLTLKINELVIVVANLKPKTMRGIISHGMLLCASLD--GD----------TK
             :*****:****:**:**:***********:**** ***:**::*:****::**** *:*******    **          :*

C.hominis    EYCEPVSVPKDAKVGELIFYNDFKGEPDVVLNTKTGKDPFAAVQPNYNVSSDLLCRFKESTMMTSAGPVFVENKNLIHGK
C.parvum     EYCEPVSVPKDAKVGELIFYNDFKGEPDVVLNTKTGKDPFAAVQPNYNVSSDLLCRFKESTMMTSAGPVFVENKNLIHGK
C.muris      -LCEPIIVPNGAKVGELITYEGFSGSPDPVLSSKAGKDAFSTVQAHYNISNNSICRYKDIIMMSSAGPLYV-NRIFAGAK
              ***: **:.******* *:.*.*.** **.:*:***.*::**.:**:*.:.:**:*: :**:****::* *: :  .*

C.hominis    LS
C.parvum     LS
C.muris      LC
             *.
```

### Figure S2

##### ***TOXOPLASMA*** (*Tg*-p43) / ***NEOSPORA*** (*Nc*-p47)

```
             β1          α1                  β2            β3
T.gondii     MELIYDSTPAALATVLLARYVQPCLLQRQPPLSLSLCFLGNEAKAYLKAQEAELPVLLAAKEGAGAEPAFPPLREPATIL
N.caninum    MELLYDSTPAGLATALVGRYVKPCLLQRQPPVPLSLCFMGTEAKAYLKVNEVRNSDTSRP--------Q---SHPPSS--
             ***:****** *** *. ***.**********:.*****:*.*******.:*.. .    .          : *::

             α2                          α3        α2?        α4
T.gondii     RRLATEAAVTPASGKSSSAGALLLGGESDVEQAQVSQWLSFCCENNFAIRGASLFQHLHSHLATRTFFCGDRLTIADFAV
N.caninum    -------HV-PSIVK-------------PRTAAQVSQWLSFCCENNFAIRGASLFQNLHSHLATRTFFCGEHLTIADFAV
                    * *:   *                **********************.*************::********

             α5         α6      α7
T.gondii     FVSCYAWMASSSQKERVLYCNLTRWMDYIQHLPGVMNAVEQLPLLSLLPPPQPVAAPPTDSAAASTQALAKQQAASGSKQ
N.caninum    FVSCYAWMASASQKERILYCNLTRWMDYIQHLPGVMNSVDQLPLLNLLPPPQTVSAAPSDAGAASAQAAAKQQAGCGTKQ
             *********:*****:*******************:*:*****.******.*:*.*:*:.***:** *****..*:**

T.gondii     GKKKSGELHVNGDCRPNGDVPKSQKKEEKGDKGAGGDHKKVSGRAKQSTPGS-SKGGAPGADSSRPVEDVTRLCMVVGQV
N.caninum    SSKKSAEPQMSGECRSND-APK--KK---GEKGAGGNDKKGSGRAKQSTAGSSSNGGAAAADSSRPVEDVTRLRMVVGQV
             ..***.* ::.*:**.*. .**   **    *:*****:.** ********.** *:***..************* ******

T.gondii     KKVWRHPEADKLFCEEVDIGEP-QVRMIASGLVPYMKAEELEGQKVIVLANMKPKNLRGFPSHGMLVCATSKDHSKCELM
N.caninum    KKVWRHPEADKLYCEEVDIGESGQVRMIASGLVPYMKPEELEGQKVVVLANMKPKNLRGFASHGMLICATSKDHSKCELM
             ************:********. **************.********:*************.*****:*************

T.gondii     RPPPDTPIGERLTFEGLQGEPDPVMNTKEGKYPFAAVQPHFYTDDNRVGMYKTHRFMTKQGPVFCDSIVGGTIS
N.caninum    RPPPDTPIGEQLTFEGLQGEPDPVMNTKEGKYPFAAVQPHFYTDENRVGMYKTHRFMTKQGPVFCDSIVGGTIS
             **********:**********************************:*****************************
```

#### D GST-like ERSs

##### *PLASMODIUM*

```
                     β1             α1                   β2          β3
P.vivax       MVESDGINIYYGK-KYPFVCRTILNAHKSLRQKENPAG-GDHSAEE-KVKFVKDEN-VEELKVEFVS-------------
P.yoelii      MIKDSKINIYYGK-NYPFLCRTVFNIYQNNIKKKTANNLNDKKIEEICVNFINDKTIVEDIKIEFVK-------------
P.berghei     MIKDSKINIYYGK-NYPFLCRTVFNIYQNNIKKKTANNLNDKKIKEICVNFINDKTVVEDIKVEFVR-------------
P.knowlesi    MIGGDKIIIYYGK-KYPFVCRTILNVHKILSQKENVEGRSNHFPND-KVKFVKDEN-VEEIKVEFVN-------------
P.falciparum  MIENTKVNIFYGNNKYPFICKSILNVYKNKNKKILEKTETTVNVDE-NIKFIKDAN-VEELKIEFLTNAPNNNNNNNNNN
P.chabaudi    MIKDSKINIYYGK-NYPFLCRTVFNIYQSNLKKKTAN-LTDQNIEEVCVNFINDKKVVEDIKVEFIR-------------
              *:  .   : *:**: :***:*::::* ::    :*              .:   ::*::* . **::*:**:

                                                         β4      α2
P.vivax       -----------------------------------KEEAQGEKCINTTDKIFAKSLDQILQTKLYHSF--RENNKKEDKQ
P.yoelii      ----------------------------NSVADESRKNVNTNNSVISSDKIFAINLDFLLKTNLYYFTSYRENINK-NII
P.berghei     ----------------------------NTVSDQTRKNTNTNNSVTSSDKIFAINLDFLLKTNLYYFTSYRENINR-NII
P.knowlesi    ----------------------------------KAEEENGDKCINTTDKIFAKTLDEILQTKLYHSF--CENNKKEDKQ
P.falciparum  NNNNNNNNNNNNNNINKGNMNHCGKNQEEEKKNLSKSNNIEKVPYICSSDKIFAKILDHLLKTDLYNIS--EKIDPKNNKV
P.chabaudi    -----------------------------SSVSEEASKNPNTINVSSSDKIFAINLDFLLKTNLYYFTCYKEIINK-NII
                                                        : ::*****  ** :*:*.**      :   : :

                  α3               α4                 α5                α6
P.vivax       TDVYVQAQYDEWVDYFRSKQLQKEVLIICDHLNRHLHLNTFICGDYLTLSDLYVYYEMHRYFNVANCYNVKYSKQFRNVN
P.yoelii      TNVFFQAQYNEWIDFLRNKDIEKNIIPICEHINKHLYLNTFLSFHYLTLSDIYIYYEMHKYFSGNITTNLKYPKQYKNIN
P.berghei     TNVFFQAQYNEWIDFLRNKDIEKNIIPICEHINKHLYLNTFLSFHYLTLSDIYIYYEMHKYFSGNITTNLKYPKQYKNIN
P.knowlesi    TDVYVQAQYDEWVDYFRNKHMQKEILIICEYVNKHLHLNTFICGEYLTLSDLYVYYEMYRYFNVANCYNVKYTKQFRNIN
P.falciparum  VNIYVQTQYEEWLDFFRSRNVNKDIVFICEHLNKHLHLNTFVVSEYLTLSDIFIYFQMYKYFHVSSTQYAKYSKMYKNIN
P.chabaudi    TNVFFQAQYNEWIDFLRNKDIEKDIMPICEHINKHLYLNTFLSSHYLTLSDIYIYYEMHKYFSGSITTNLKYPKQYKNIN
              .:::.*:**:**:*::*.:.::*::: **:.::*:**:****:  .******:::*::*::**        **.* ::*:*

                α7          α8
P.vivax       RWFKLINCLVN---YSDAELSTMLKKEVETNQVDITK------------DANTMQLLKQKIVSTSYV
P.yoelii      RWFKLIKSLLHDHITHDAELVQNLKIKEKIFNTTNYGNKKSGNNEDKSSTHPNKSNTKVGGTSLSSYE
P.berghei     RWFRLIKALLHDHVATDAELIQNLKVKEKIFNTAGSG-KRLGNNGNKYSAHSDHTNTKGGATSLSSYE
P.knowlesi    RWFKLINCLVN---YPDEELSTMVKKELEKNQVNITK------------DANSKQLLKQKTVCTSYG
P.falciparum  RWHKLIDSLIY---FEDAEWKKNLKCLICDNKMNFDDRKDNLKNNNKSMNNTDDVKC-KKKTVTTSYS
P.chabaudi    RWFKLIKSLIHDHISTDAELIQHLKIKERMVSTGGAG-RRPGNNADKSSGD-------HGKGSAASYE
              **.:**..*:      * *    :*       .                 .            :**
```

### Figure S2

#### THEILERIA

```
                       β1         α1             β2          β3  β4    α2
T.annulata      MYNKGSEQLVVNYSVQNPPYGAVILSSISKYLQKSDSSKVVFNLDNKLQNNEFSLNGDKFNEVDLLKNLCSVLPYCDKLL
T.orientalis    MKASNPCSLVVNYSVQNPPYGAVILSCISKYLQKSESSPVEFNLDNKLQNNEFSLNGNNLNEVELAKKLTTLLPYCDKLL
T.parva         MYSKGSEQLVVNYSVQNPPYGAVILSSISKYLQKSDSSKVVFNLDNKLQNNEFSLNGDKFNEVELLKNLCSVLPHCDKLL
                *   ... .*****************.********:** * ****************:::***:* *:* ::**:*****

                        α3                 α4                α5          α6
T.annulata      DVVNNSELEYWFKVLLSEKSKLAENKGNFTENDSYLENMNKHLAQRTYLIGHRLSLSDILHFSLLRRTCNVNSLKNKYPH
T.orientalis    DVPEYSELDYWLKVLLLEREKHLETKKAFFENVEYFEQMNNHLAHRTYLIGYRLSLSDVVHFSFLRSNCNVNSLKNKYPH
T.parva         DVVNHSELEYWFKVLLSERAKLTENKGTLAENDSYLENMNNHLAQRTYLIGHRLSLSDILHFSLLRRTCNVNSLKNKYPH
                ** : ***:**:**** *: *  *.*   : ** .*:*:**:***:******:******::***:** .************

                  α7
T.annulata      LARWYNFISNVP-GTSGCLNGYETPSSATYKKPFTKQEQQDNSYK
T.orientalis    LVRWYNFISNVPVGTYECLKKYDAPARVQYKKPFTKQEQQDNSYK
T.parva         LVRWYNFISNVP-GTSGCLKGYDTPSSATYKKPFTKQEQQDNSYK
                *.********** **  **: *::*: . ****************
```

#### BABESIA

```
                       β1         α1             β2          β3  β4    α2
B.microti       -----MYEFNVLYHPTCTILG---IDAFAN-LTRKEGIKVTLICDKGANVNSYTIKSRDSILTESFSCELSALDYLSGWS
B.bigemina      MAETTRDAVTVVHAPRETPYGAIVLAAIAKRFSKANESPIAFEAGKAIEPGTYLLEK-AKLQEPQLLRELCLRIPSCKRM
B.bovis         MDNGSKCAVTVQHAPKDVPFAAIVIDAIAKQLSRSDDDVVKLEQNKAITPGTYVLDK-VSMQISQLVRELCLRVSTCKRM
                        ..* : *  .  .   : *:*: ::: :    : :  .*.   .:* :..  .:   .:  **.     .

                         α3               α4                α5          α6
B.microti       KNFQPQENIEIWAEILTNCSKDETLM--------AALDKYLSLKTFLSGHSMSCIDIAMFSIL-------KPDPPYPNVT
B.bigemina      LYIDSSASLDQWINYVTLHSATPKTPVADLWQYYNKMNTHLATRTFLIGYRMTVVDVLQFAALSNAKAVCKELAKVPHVQ
B.bovis         LYIDSTASLDKWLSYISEGAGMHKT-EEQLWQHYTKINQHLTMRTFLVGYRMTAVDVLQCSLLANSQGIRKRLTELKHLE
                  ::.  .:: * . ::  :    .               :: :*: :*** *: *: :*:    : *       *  .   ::

                  α7
B.microti       RWHRFIAHLIGPTISTN-----D-------KFGNQGNVRKAQVDNSYR
B.bigemina      RWYNYIIAIPGVQQAVS-----DIQRKNVPQVEKKPFTKEQQIDNSYK
B.bovis         RWFNYIMAIPGVNSNPGTATTNRAENKVFTKNDKKPFTKAEQFENSYK
                **..:*  : *      .                :  ::  .:  *.:***:
```

#### TOXOPLASMA / NEOSPORA

```
T.gondii        MHSRGTPGQKRRGRRHRGVANAENRLYAAVSRHNFSLNSLLTFELGVSVWPEQELLRNPLLVASPTSTLRPARKRRVESL
N.caninum       MLRVRY------------SPQHPPLATLCVSVL---------------LERGLAACVPDNKLQGS--------
                *                          **                 :.:.
                 β1                    α1                                β2

                               α1                        β2         α2      β4
T.gondii        TRCRDTAVTFDGARAFSFFVFFLRFLACLEEPRCCAVQKTESGSEEVFTDIAAAKLLCTLSSKTRLLYPAAAPAPSFADA
N.caninum       ---------------------------------LLIVQKSDNAPEEFTTDVAAAKILCALSPKTLLLYPAAVPSPSFEDA
                                                 ***::...**.  **:****:**:**.** ******.*:*** **
                                                  β3        β4

                        α3                   α4                     α5             α6
T.gondii        SETACIDTMLSLATSLQV-SSLSPKQLGSLSSHLQLRTFLCGFHLSLADLAVYTQLRRHANNEQQAGAPPQGWKDKFVHV
N.caninum       CATACIDAMLSLAATLQAASSLSPKQLGSLSSHLQLRTFLCGYHLTLADLAVYAQLRRLACTEQHAGAPPAGWKDKFVHV
                . *****:*****::**. **********************:**:*******:**** *  .**:***** *********

                  α7        α8
T.gondii        SRWYAFIHGQPQISNVVAVALRKAGANAKGAEASKNE-GKKREGRANSSGASSPPSADKRAAAAQASYE
N.caninum       SRWYSFLHGQQSTANAVGVALRKAGSTATRAESQKNEDGKKKAERANA-NPSSPTAGDKKGTASHASYE
                ****:*:***  . :*.*.*******:.*. **:.*** ***:  ***: ..***.:.**:.:*::****
```

**Figure S2**

### E GST-like MRSs

#### *PLASMODIUM*

```
                 β1        α1           β2                        β3   β4      α2
P.vivax      ------MILYPHKYFPN-TVKCMLAAHIYSFQVELCDDFFYTRSFDIEKSLVVNKKPLLVYEDNYVSSTKAICFLFHRLR
P.yoelii     ------MILYPNKYLLSGTLKCMLAAHIYKLPVELSDDFFFTRTFEVDKSLIVNKKPLLIYENKFISSTNAICFLFHRLK
P.berghei    ------MILYPNKYLLSGTLKCMLAAHIYKLPLELSDDFFFTRTFEIDKSLIVNKKPLLIYENKFVSSTNAICFLFHRLK
P.knowlesi   ------MILYPHKYLPN-TVKCMLAAHIYSFPVELSDDFFYTRSFDIEKSLVVNKKPLLVYENNYVSSTKAICFLFHRLR
P.berghei    MMKPLNMILYPHKFFLN-TVKCMLAAHIYNFNVELSEDFLYTRTFDIEKSLIINKKPLLIYENNYVSSTKAICFLFHRLR
P.chabaudi   ------MILYPNKYLLSGTLKCMLAAHIYKLPVELSDDFYFTRTFEIEKSLVVNKRPLLIYENKFVSSTNAICFLLHRLK
                   *****:*::  . *:*********.:  :**.:**  :**:*:::***::**:***:**::::***:*****:***:

                             α3            α4
P.vivax      NVE-SKKCVDEKLRLYMSWVEWSDLLEKHIEVLNKKKIIDSLDELESYLKENP--NKNFICSL----------GEATQGD
P.yoelii     NKGINTKGLDDKLRLYLTWIEWSDILEKNIERLNKKKIIESLDLLESYLSENK--NKKFICDEVDQIKREDECSETDQNQ
P.berghei    NKGINTKGLDDKLRLYLTWIEWSDILEKNIEHLNKKKIIESLDLLESYLSENN--NKKFICDEVDQIKREDGCNGANQNP
P.knowlesi   NAE-SKKCVDKKLQMYMSWVEWSDMLEKHIEVLNKKKIIDSLDELESYLKENQ--NKNFICSC----------EQDSQGD
P.berghei    NK--SKKPFDENLSYYLGWLEWSDLLEKHIEVLNKKNIIESLDELELYLKENKNKNKNFI--------------NNDDEK
P.chabaudi   NKGINAKSFDDKLRLYLAWIEWSDILEKNIERLNKKKIIESLDVLESYLSENN--NKEFICDDAEQIKREDGCDETDQSQ
             *    . * .*.:*  *: *:****:***:**  ****:**:***  **  **.**    **:**                  :

                        α5                     α7                  α8
P.vivax      SND--KLTLADIFVYTSLTHCNIEVCRESWAHISNYIEKINQLEDVQKILKDVDRIYKYKNIYNLFISKIHEKNINTYLK
P.yoelii     KNDICGITLADIYVYTSLKYSNIVICNESWSHINNYIERIHNLEDVQKVVETVEKVHLKLKNIYSLFISKVCEKNINDHVK
P.berghei    KNGICGITLADIYVYTSLKYSNIVICNESWSHINSYVERINNLEDVQKVVEAVEKVHLKLKNIYSLFISKTCEKNINDHVK
P.knowlesi   GND--KITLADIFVYISLTHCNIEICGESWIHINEYIQKINQLEYVQKILKDVEHIYKYKNIYNLFISKIHEKNINSYLK
P.berghei    DEN--KLSLADIFVYTSLKHSCIEICKESWVHINEYIEKINNLQEIQNVIKNVEEIYKYKNIYSVFISKIHDKNVNEYLK
P.chabaudi   ENGTCGISLADIYVYTSLKYSNIVICNESWSHISSYVERINNLEDVQKVIETVEKVHLKLKNIYSLFISKICEKNINDHVK
              :.   ::****:** **.:.  *  :* *** **..*:::*::*: :*::::  *:.::* ****.:****   :**:* ::*

P.vivax      EEKFYI
P.yoelii     NDNFYI
P.berghei    NDNFYI
P.knowlesi   KEKFYI
P.berghei    NETFYI
P.chabaudi   NDNFYV
             ::.**:
```

#### *THEILERIA*

```
               β1        α1             β2
T.annulata     MEHHLLILNDSNESHLCRFLLKTLEPSFPLVVLYGPKVPDKLSDLYIESSELEHLKDSTTDSNNYKYSVLYKELNKSNSL
T.orientalis   MEYHLLILNKSNESLLCRNFIKIWASSAPVVSLFSSKIPEDLSDIHIDSDDHDTFKDSTSEANSYKYQQLYNELNRLCTL
T.parva        MAHHLLILNDSNESQLCRFLLKTWEPSFPLVVLYGPKIPDKLSDLYIESSELDHLKDSATDSNNYKYTVLYKELNKLNSL
               * :******.**** *** ::*   .* *:* *:..*:*:.***::*:*.: : :***::::*.***  **:***:  :*

                                                 α2                        α3
T.annulata     THQEFEELYNNLVKYLKTVLFRPVLLLFKKSNNFLPNVLFNFKPILLYLSPLYNLHKVDGQKKQYREDKNLLYWYLEFDY
T.orientalis   SQQDFSIIHTNLFKYLKRALFRPLLLLFKKDNEFVPNIVFNLKPILLYYYPS--LRKVSDTQ---KEDKELLYWYTEFHS
T.parva        TQQEFEELYNNLLKHLKTVLFRPLLLLFKKTNNFLPNVLFNFKPILLHFYPLYNFPKVDGSNNTYREDKNLLYWYLEFDY
               ::*:*. ::.**.*:**  .****:****** *:*:**::**:*****:   *    : **.. :   :***:***** **.

                      α4                α5                  α7
T.annulata     NLNNIGTNNRLLDLLDELENTIEEYNLDNKFVQSILSLLNMYVDRILFRNISLTKYTKVLKYSKQYSTLLNT--------
T.orientalis   NFNYVGTNRKLVELLDDLEENLSRYNLESPLTQLLLSTLNMYIDGLLFRNIALSKYTNILRQATHHKHLLKNSLLNKFNY
T.parva        NLNSIGTNNRLLELLDELENTIEEYNLDNKFVQSLLSLLNMYFDRLLFRNISLTKYTKVLKYSKLYYSLLNT--------
               *:* :***.:*::***:**:.:..***:. :.* :** ****.* :*****:*:***::*: :. :  **:.

T.annulata     ----HLSSTGTYTHA--------KGS-------TTE-AQETS-------------------SKFTEKGEVGHE-----EL
T.orientalis   QGTATVSATGILSPSIPTSLSGFKGSDKANGASKGK-SESAKSPLKGSGKGAESSSVYNMGSGKEGKEDAEKEWEPGLER
T.parva        ----HLSSTSTYTPP--------KGS-------STPSTQDTG-------------------SKMSENPEVGPE-----EL
                    :*:*.  : .          ***        .   ::.:                    *    : :.  *     *

T.annulata     DKSSESADKLNLVDYK
T.orientalis   DDDLEEDEKLSLNDYK
T.parva        DKSSGAPDKLSLVDYR
               *..    :**.* **:
```

Figure S2

#### BABESIA

```
                β1          α1           β2
B.bigemina     MQ--LLVLDSSIDSLVTWSAAECVCDSVPTVVLASISTESDLRSNKCRNEVHALKCLQELPLLCSKLESQDAHSVKLFTE
B.bovis        MQ--ILILDQSLESLAAWLTAEFLRNSSPLALLSSVTRDTNWNPSASFEEHNGYKCVNTLSLSCENFEQNDGDFHGFLES
B.microti      MQYTLLLDPSCYHSFACYLASAQFNFPAKIEIAADVNIE-----NRLVSEEYNLSKFKDKKLQIATLDT----------
               **  :*:  .. .*:. : ::  .  .    : :.:. :      .    .*     . .:    *    .::

                                                  β3       β4     α2
B.bigemina     HIGKHTSQPSEDVLKH---L-LKTLNEVPVRPLLFAVDSSVADVLPIFGCNAVLKYYVEGQGVCGKSSCDLYTPTYEQSA
B.bovis        IFGLNPEACDTSTVKSTKAIIVQAQKDILVRPLLLIVNRNNLEVSPIFGSDSILRYYLENQTSTINGNNILLNSEQVKNH
B.microti      ------------------------------KPVLIVNIGN--DLIKIKGIRAILLYIYRY------------TIKDIDNF
                                             :*:*:    .  ::  * *  ::* *  .             .    ..

                  α3                 α4                       α5                      α7
B.bigemina     FDWFSWTSQLRLLCKKNASVDLRALLSQFDAYLHGQQECDIKVGIGSVVITLCLLWYETKLEFLGCNLDMFSHIRQYISA
B.bovis        FDWKSWSSTLRTVCRKNNSDLARGLLSSLNAFLTSNIDLICYDIIGSAIITLSILWFNGKIEALGLRLDMFPSIPPFIHN
B.microti      YNIYHYFD----------------LLERLEALLTGVNKLHIDNILQK---------LDDHLSTSDYLLDKITPVDIYLYS
               ::   : .                **. ::* *  .  .      : .          : ::.   .  ** :. :  ::

                           α8
B.bigemina     VLTATANVERYRCLHTFYRLHGTCAAPERSLASSRELE--------SLCRDFIDR-KFYV
B.bovis        VLTKTSSLNKARCLHLFYRSMETSRSHVNGNGALKSLS--------DVSRDLIGK-KYYI
B.microti      MLSAMHNFH-WKNLSKFHSNLHMFANHLSKFKLTQQFYDYSTQPYDNLSVDIKKCKNYYI
               :*:    ...  : *  *:                :.:          .:. *:    ::*:
```

#### TOXOPLASMA / NEOSPORA

```
                                   β1          α1         β2
T.gondii       MSTMTGSSPKSAASSTPLRLVAGSAVPRGDVAACIAVGHALNIPVVLVSKDADKAEQAEPSSSAPAPPPQAPVASSASEE
N.caninum      ---MTDSA-PSGASSSPLRLLAGCTVPRAEVAACLAVANALKVPVVLSSKDAAES-KARPSQASLAPTPHAPASDPTVDE
                  **.*:  *.***:****:**.:***.:****:**.:**::**** ****  :: :*.**.:: **.*:**.:..: :*

                β3    β4      α2                          α3
T.gondii       DKKDLCVLLTREGSVVSAYAICLYLLSLARKRLAETKTAAP--EKRLESREKLSEAYDGDAWQAEDALAWAAFKLRGALR
N.caninum      DKKDACVLVTPDGSVVSAYAVCLYFLSVARKRRRQAEKAAPQQEKRQEGEAAPARDAERDAWQAEDALAWAAFKLRGALR
               **** ***:* :********:***:**:****   :::.***  *** *..    :.   : ********************

                    α4                            α5                       α7
T.gondii       GVRKDLLLQQLNLLEERIAG---SSDCAFPALSALLSTSAPSPFVAPAMFVFCMLRYAHPTSAFSDWPALSAFLGSLGAS
N.caninum      GVRKDRLLQQLNSLEERLSGDPQTSPAAFPSVSSLLAA-SPSLFVAPAMFMFCLLRYAHPTSSFADWPSLAKFLQALGSS
               ***** ****** ****::*    :* .***::*:**:: :** *******:**:********:*:***:*: ** :**:*

T.gondii       PAAAATLKAVPLPARSRGFPEMSQALLASAGACAVAQRQREKFYI
N.caninum      PAVAATLHAVPLPSRACGARALTRALLSSSA-SPQTEKTRERFYI
               **.****:*****:*: *   :::***:*:. .. ::: **:***
```

### Figure S2

### F GST-like QRSs

#### *PLASMODIUM*

```
                 β1            α1            β2      β3      β4      α2
P.vivax         MENVEKIYIRNKAELSDNFFYFYNWVKLNFNTCLIISENSSLRSSVLLELLNGKKLVLSEINVCEALFRLACGGI-----
P.yoelii        MEHIEKIYIKNKSDLGNEFFYFYNWVKEKFNISLVISESNSLRHNIVLELSDGNNLSLNELNSCEAIFHLISNDK-----
P.berghei       MEHVEKIYIKNTSDLGNEFFYFYNWIKEKFNISLIINESNSLRHNIVLELSEGNNISLNELNSCEVIFHIIRNSH-----
P.knowlesi      MENVEKIYIRNKADLSDNFFYFYNWVKLNFNTCLIISENSSLRSSILLELLNGKKLVLSEINACEALFRLTCGGV-----
P.falciparum    ---MEKIYSKNKLDYLSIIFNFYSWYKRHFKGTLLINENSNIKNTTLIELLGNKKLLLNEINACEAILKLNEHHENYDSS
P.chabaudi      MEPIEKIYIKNKSDLGNEFFYFYNWIKEKFNTSLIISESSSLRHNIVLELPEGYKLPLNELNSCEVIFRIICNGY-----
                   :**** :*. :   . :* **.* * :*:  *:*.*...:: . ::**  . :: *.*:* **.::::

P.vivax         -----------SGE---------------------------------------SSGGVSG-----------EISDVSCHH
P.yoelii        -----------NSKNS-------KN-N---------------------QNNKHIWNNEFE-----------SEQDVEENP
P.berghei       -----------NKK---------------------------------------DNWGNEFE-----------NEQDVEENP
P.knowlesi      -----------SNAVG-------GTVS--------------------CTVNCLSNGPVSG-----------AQRDDFANR
P.falciparum    SGTCNVNNINQPNNNNNNKENINDNNNNNKENINDNNNNNKENNINPCSTNSFNINSFSTNNYNNFNNPCCSFSCNIYDY
P.chabaudi      -----------NKN---------------------------------------NIWGNEFE-----------NEKDKEENP
                                                                       .                        .

                       α3             α4                          α5
P.vivax         RIDALYAEELKFFLKGRLASDKPVDEQFLERVEAEMALQ--KGFCKER---LSFLDFLLFSIFKKAKVDQVEG--NKYPH
P.yoelii        -FRISQIEELKNFLKNILLQKKKLENNFYERIEKEMMNKNNNIFSKDK---LRYIDFLLFCILKKNKIEKIN---NKYEH
P.berghei       -FKIVQIEELKQFLKNILLQKKKLENDFFEKIENEMIKKNNNIFAKDK---LRYVDFLLFCILKKNKIEKIN---PKYEN
P.knowlesi      CRDTLRGEELKFYLKEKLASEKPVDEQFLQRVEAEMALQ--KVFCKDK---LSFLDFLLFSIFKKGKVEKLKDPLGRYPH
P.falciparum    -YDILKIEELKDFLKDFILRNKKINKEFFDTIEEDMTMWKKIFFFGQKKRTPEFLDFLLFYILKKENIHFFEE-ETRYIN
P.chabaudi      -FKIVQIEELKHFLKDTLLQKKKLDDNFFEKIENEMICKN-TIFSKDK---LKYTDFLLFYILKQNKIEKIN---PKYEN
                         **** :**  :  .* ::.:* : :* :*       *  ::      : ***** *:*: ::. .:    :* :

                 α7
P.vivax         LGAWLSKNN---FP--IKEEDMNSFAS
P.yoelii        LNNWYLQNN---IS--VKEESMNSFTN
P.berghei       LNNWFLKNN---IS--IKEESMSSFTN
P.knowlesi      LSAWFSKNN---FT--IKEEDMNSFAS
P.falciparum    LHNWFLKNKIYDISNDKKEDNTPSACN
P.chabaudi      LNNWFLKNN---IS--IKEESMNSFTN
                *  *  :*:   :.   **:.  *  .
```

#### *TOXOPLASMA / NEOSPORA*

```
                     β1               α1           β2              β3
T.gondii        MALSPGAVLRITRLASLPPLSTVTDVVLAYIEQNLKSAGDFTFSVQPDTSSTPAAPFRIAVETKKGEQPSPIAAGEALSD
N.caninum       MALSPGAVIRVTRLSSLPPLSTVTDVVLSYLEQHLKSAGGFTFSVYPDTSSSPATPFRIAVETKKGEAPRPIAGGEALSD
                ********:*:***:*************:*:**:*****.***** *****:**:************ * ***.******

                     α2                          α3                           α4
T.gondii        YAAARCIARLAASTGDPFASLLKASGAPGGVEAAVDVCEQWAAFYGVSGKAPEASDNPGLSACIDFFARRFSTSSSFLFG
N.caninum       YAAARCVARLAASTGGVFACLLNENGATESADGAVDACEQWAAFCGVPGKTPEAAEHPGLTACIEFFTRRFSTSSAFLCN
                ******:********. **.**:  .**.  ..:.***.*******  **.**:***::***:***:**:*******:**  .

                          α5            α7
T.gondii        TPQPSLADMTLFALLKAHPAAQTALSSALRTSETAAKHFSSWSACVAKAIPGAGALSGAGAKGTDRAKGKSGEMVG-QGA
N.caninum       TPNPSLADMALFALLKTQPAAHTALLSALRTSGEAAKHFSAWSSSIAKALPGATTLSGAGVPD-GSAQGNRGEISGATGA
                **:******:******::***:*** ******  ******:**:.:***:***  :*****. . .  *:*: **: *  **

T.gondii        GVASEAVDGPDATTS
N.caninum       GASPESTDGPDATAS
                *.:.*:.******:*
```

Figure S2

#### G N-terminal extensions of YRSs

```
P.yoelii        ----------------------------------------------------------------------METINIKKED-NE-----------AQVKISQEIH-DENVEKRYNEIMSITSEC
P.berghei       ----------------------------------------------------------------------METINIKKED-NE-----------AEVKISQEIH-DENVEKRYNEIMSIASEC
P.knowlesi      ----------------------------------------------------------------------MESDGVKREE-LQGATGEAAGLVEAQETAPQENPPQEDVSRRMTEILSIASEC
P.falciparum    ----------------------------------------------------------------------METTDTKREE-QE-----------IEEKKAQEESKIEDVDKILNDILSISSEC
P.chabaudi      ----------------------------------------------------------------------METINIKKED-NE-----------AQVKIPQEIH-DEEVEKRYNDIMSITSEC
P.vivax         ----------------------------------------------------------------------MEGEGVKREE-LEGATGEAAPQAASQEAASQEATPQEDVSAKLADILSVASEC
T.annulata      -----------------MSERFISPFHGGIAVNYS------GNVVN-----------VKNPNLEFHHDSCSIKCGLREEL---------DRYFRGS----RSQMDLTAEEKVKICLSLSEECI
T.orientalis    MLISIKNCNLSI-RLKNMSERFLSPFHGGIAVNYS------GNVAN-----------IKNPKLEFYPGSCSVKHGLRKEI---------DNFFHKS----SFHGNQSLEERVNTCLSMAEECI
T.parva         MSIYVKKSSLCIFRFKKMSERFISPFHGGIAVNYS------GNVVN-----------VKNPNLEFYNDTCSIKYGLREEL---------DRHFRGS----RPQMDVSVEEKLKICLSLSEECI
B.microti       ML---------------RFRMALSQFTGGL-YKGEIHNPLYGSKIK-----------LKQTICAFTPSQFYLEGGIRKNI---------DKYFEGK----L-STKLSVKERLDLCLQIGSECI
B.bigemina      -----------------MSRVFASPLHGGLEFDGE------EECYK------------TSTGFEFHPGSCAAKNSIRERL---------NAHFAPS----LAEPRIPVAEALERACKLSVECI
B.bovis         -----------------MMRSFVSPLHGGLDLEGT------ERSHR------------TRTDFEFYPKSCAIKNGLREEL---------ENHFRGM----QIESQLSLKDAIDICNQMSVECI
```

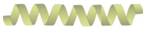

```
T.gondii        ---------------------MASPFTGGLSVW-DDSLLT-SHVASVSTAASTS-SPPA---NEASNATPATKRGEKEKKTSGGVTTLYDSIVNGAQWPQPRPSSLPLDERVQLCLSIAQECI
N.caninum       ---------------------MASPFTGGLSVW-DERLVS-SQLAALSTTASPSPAPPAPANQEANSAAVSEKQGGKDKKSSGFV-TLYDAVVNGDKWPAPRPSSLSLEEKYQLCLSVAQECI
                                       * : **:                                          : . :..          :  .             .  :  . . .:. ***
```

**Figure S2. Sequence alignment of GST-like domains present in *Apicomplexa* EF1 and multi-synthetase complexes.** Protein alignments were performed with M-Coffee (20). The corresponding EupathDB annotations are given in Table S1. The complete sequences of AIMPs are shown with their EMAPII-like domain highlighted in gray. For EF1 subunits and aaRSs, alignments were performed on the full-length protein sequences, but only the GST-like domains involved in protein-protein interactions are shown. The $\alpha$2, $\alpha$3 (interface 1 or 1') and $\alpha$7 (interface 2, especially the RW residues) helices are highlighted in bold. The N-capping box ($\varphi$-S/T-X-X-D-$\varphi$) is shown in blue in the central helix $\alpha$5, when present in the sequence. The GST-like domain of EF1$\beta$ is reduced to only 3 $\alpha$-helices including $\alpha$5 and $\alpha$7 which are sufficient to form an interface 2 with EF1$\gamma$. The strains used to run AlphaFold modelling are indicated in bold and the corresponding secondary structures are displayed above the sequences or below for N. caninum proteins.

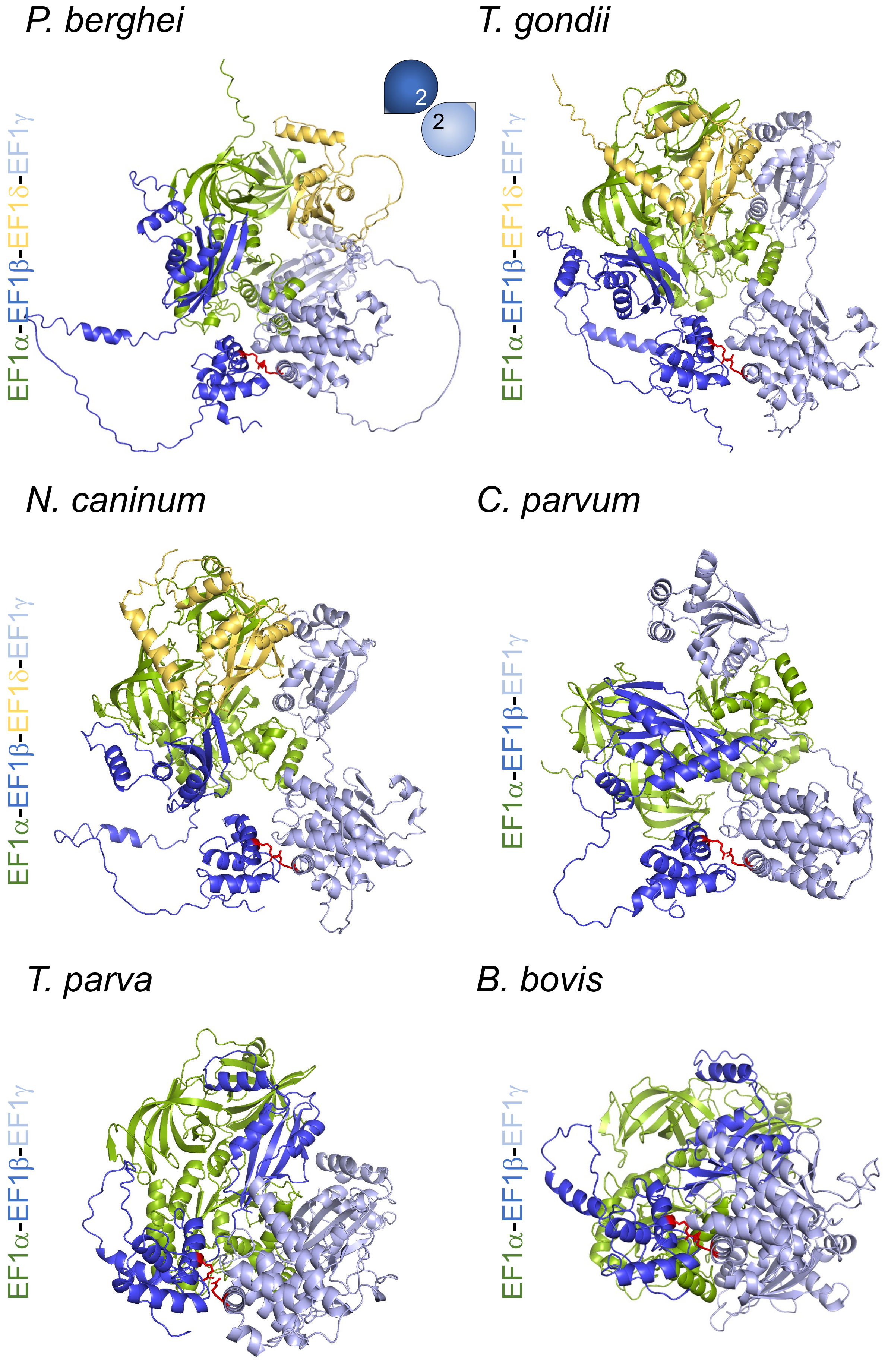
P. berghei
EF1α-EF1β-EF1δ-EF1γ
2
2
T. gondii
EF1α-EF1β-EF1δ-EF1γ
N. caninum
EF1α-EF1β-EF1δ-EF1γ
C. parvum
EF1α-EF1β-EF1γ
T. parva
EF1α-EF1β-EF1γ
B. bovis
EF1α-EF1β-EF1γ

**Figure S3**

**Figure S3. Similarities between EF1 complexes in the *Apicomplexa* phylum.** Molecules are colored in blue for EF1$\beta$, light blue for EF1$\gamma$, green for EF1$\alpha$, and yellow for EF1$\delta$. All models are shown in "cartoon". GST-like domains are schematized with drop shapes, with the same color code. *Plasmodia*, *Toxoplasma* and *Neospora* complexes are tetramers while EF1 is a trimer (EF1$\beta$, EF1$\gamma$, and EF1$\alpha$) in *Babesia*, *Theileria* and *Cryptosporidum*, due to the impossibility to identify subunit EF1$\delta$ in the corresponding genomes. All EF1$\beta$-EF1$\gamma$ heterodimers involve interface 2 shown in red; the parameters for each AlphaFold model are shown in Figure S4.

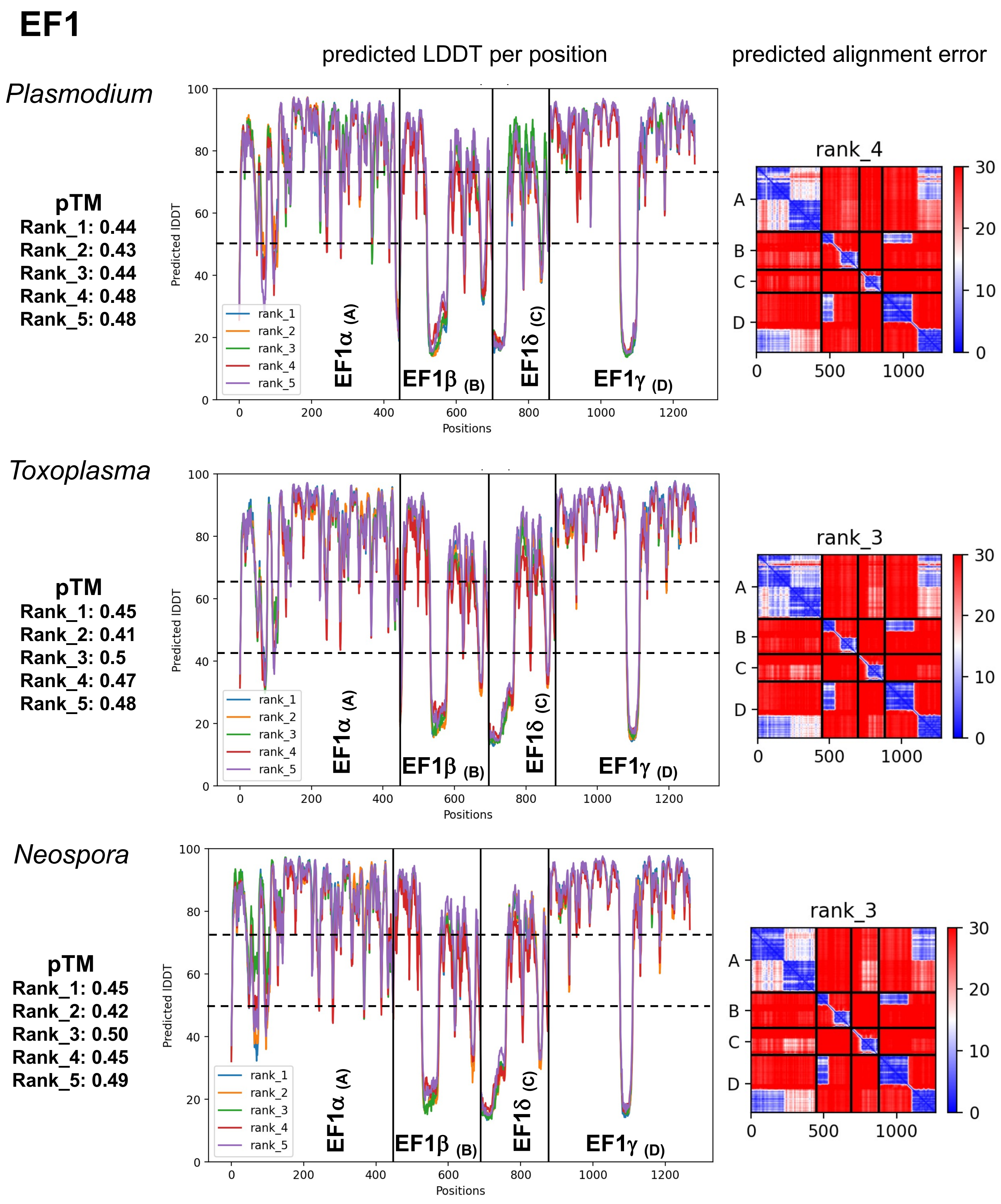
EF1
predicted LDDT per position
predicted alignment error
Plasmodium
pTM
Rank_1: 0.44
Rank_2: 0.43
Rank_3: 0.44
Rank_4: 0.48
Rank_5: 0.48
Toxoplasma
pTM
Rank_1: 0.45
Rank_2: 0.41
Rank_3: 0.5
Rank_4: 0.47
Rank_5: 0.48
Neospora
pTM
Rank_1: 0.45
Rank_2: 0.42
Rank_3: 0.50
Rank_4: 0.45
Rank_5: 0.49
Predicted lDDT
Positions
rank_1
rank_2
rank_3
rank_4
rank_5
EF1α (A)
EF1β (B)
EF1δ (C)
EF1γ (D)
rank_4
rank_3
rank_3
A
B
C
D

Figure S4

# EF1

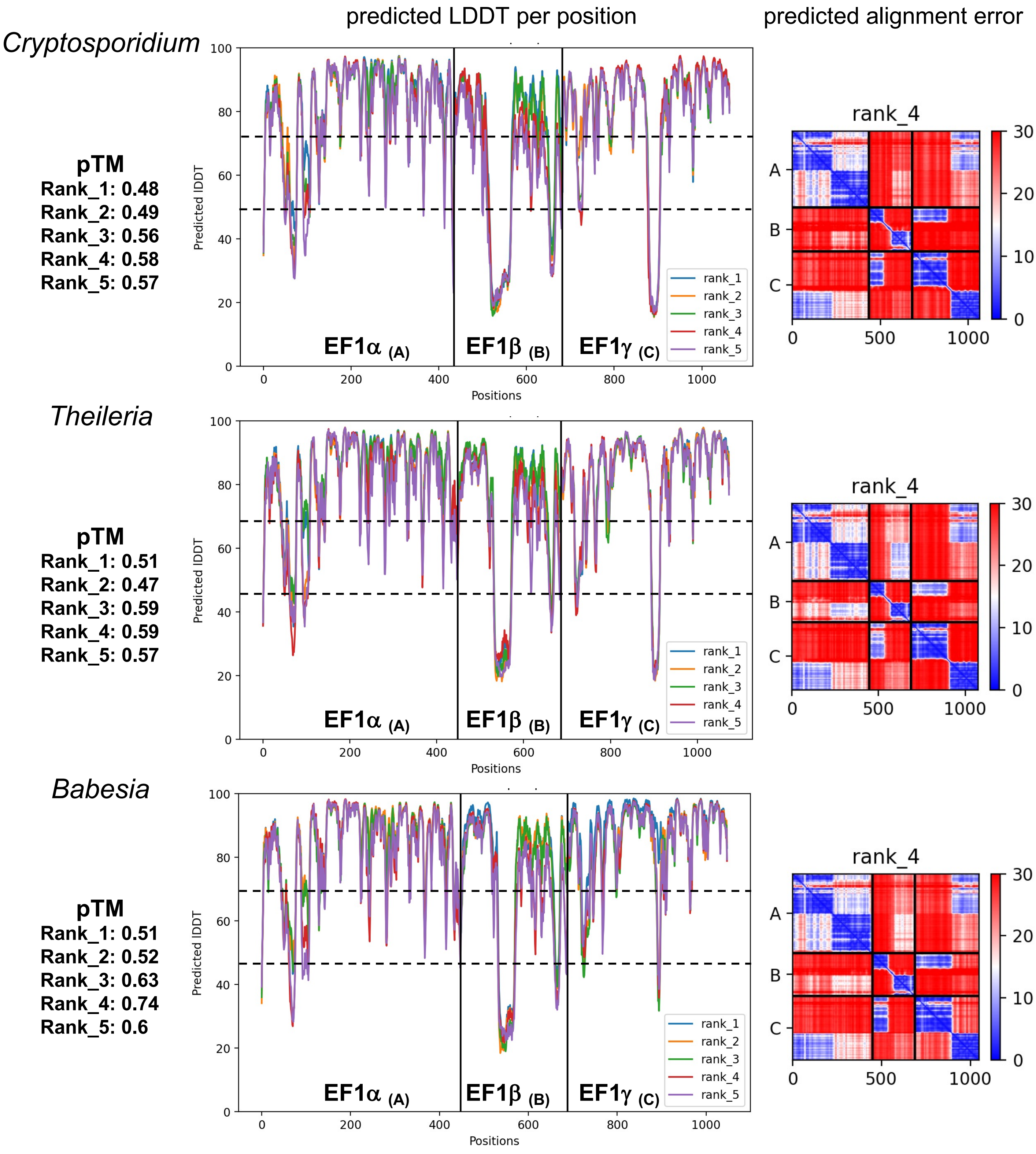


Figure S4

**Figure S4. Confidence metrics of the ColabFold predictions for EF1 GST-like backbones.** Models were ranked by pTM (predicted Template Modeling) scores, which measures the overall topological accuracy of the predictions. Predictions with pTM scores higher than 0.5 and showing the known GST-like interfaces 1, 1' and 2 were considered relevant and are highlighted in bold. The central plots show the pLDDT (predicted Local Distance Difference Test) scores per position. Regions predicted with high confidence are identified by pLDDT > 70, while disordered regions are identified by pLDDT < 50 and are presented as long filaments (not shown in graphical representations). The PAE (predicted aligned error) plot on the left indicate that the relative positions of domains are well predicted (blue).

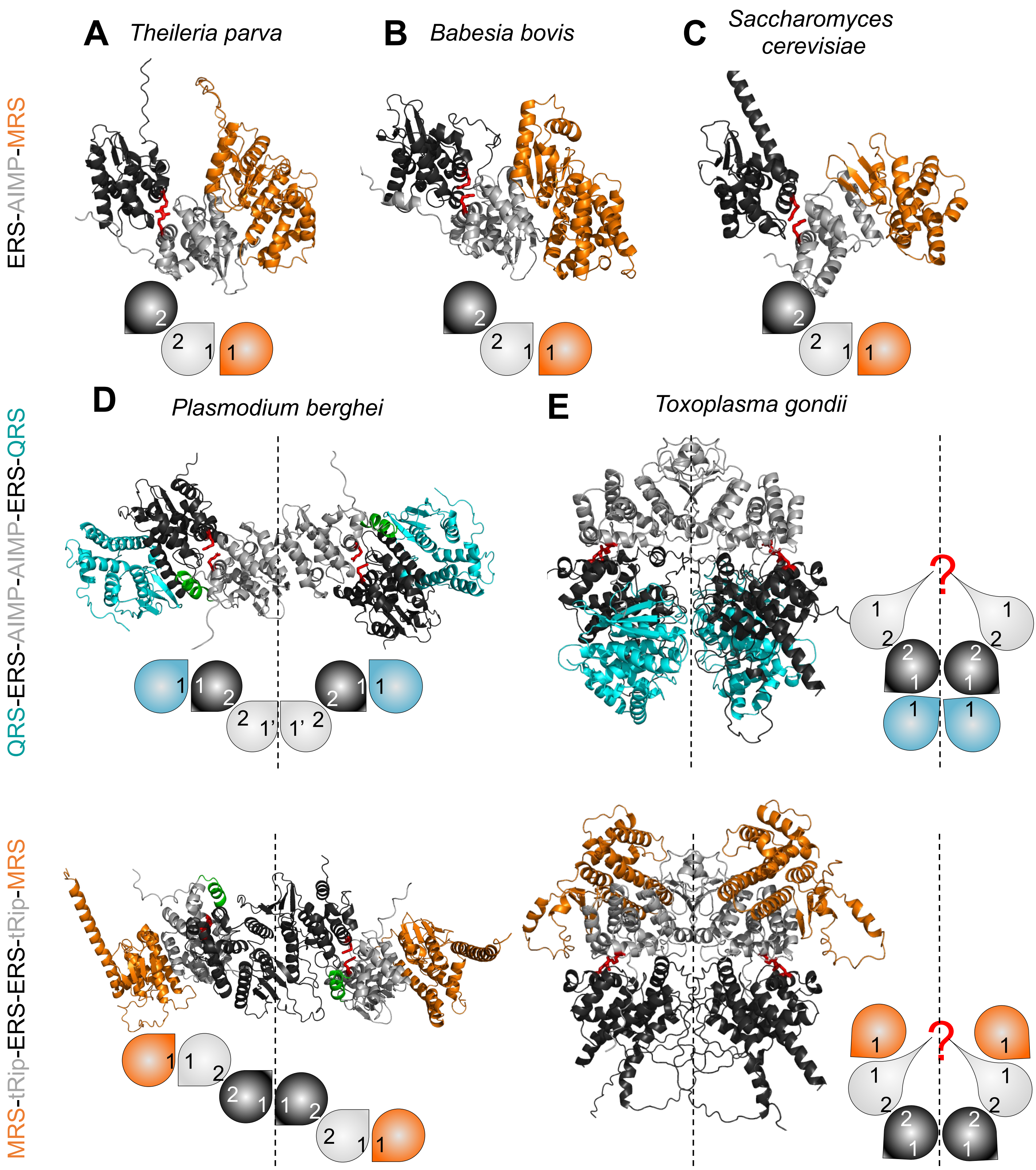

A Theileria parva
B Babesia bovis
C Saccharomyces cerevisiae
ERS-AIMP-MRS
D Plasmodium berghei
E Toxoplasma gondii
QRS-ERS-AIMP-AIMP-ERS-QRS
MRS-tRip-ERS-ERS-tRip-MRS


**Figure S5**

**Figure S5. Diversity of multi-synthetase complexes in the *Apicomplexa* phylum.** All complexes are shown in "cartoon" with the same color codes and schematized below with drop shapes colored grey for AIMP, black for ERS, blue for QRS, and orange for MRS, all displaying interfaces 1/1' and 2. In *Theileria* (**A**) and *Babesia* (**B**), MSCs are organized into AIMP-ERS-MRS trimers similar to the Arc1p-ERS-MRS trimer determined for *S. cerevisiae* MSC (**C**) (25,46). (**D**) The M- and Q- complexes of *Plasmodium* were determined experimentally (14,16). They both involve equivalent GST-like domains but organize differently. In the Q-complex, symmetry is imposed by the tRip homodimer (interface 1',(13)) that binds ERS via interface 2, which in turn binds QRS (interface 1). Alternatively, the M-complex is organized around the ERS homodimer. Accordingly, the tRip transmembrane helices $\alpha 8$ (green) are equidistant and similarly oriented. (**E**) Two *Toxoplasma* MSCs have been modeled to mimic *Plasmodium* Q- and M-MSCs. Both models show homodimerization of *Tg*-p43 through an unknown interface (?). This homomeric interaction, if biologically relevant, would allow the formation of a unique MSC (dimer of tetramers, Figure 4C). Moreover, the transmembrane helix $\alpha 8$ is not predicted for *Tg*-p43.

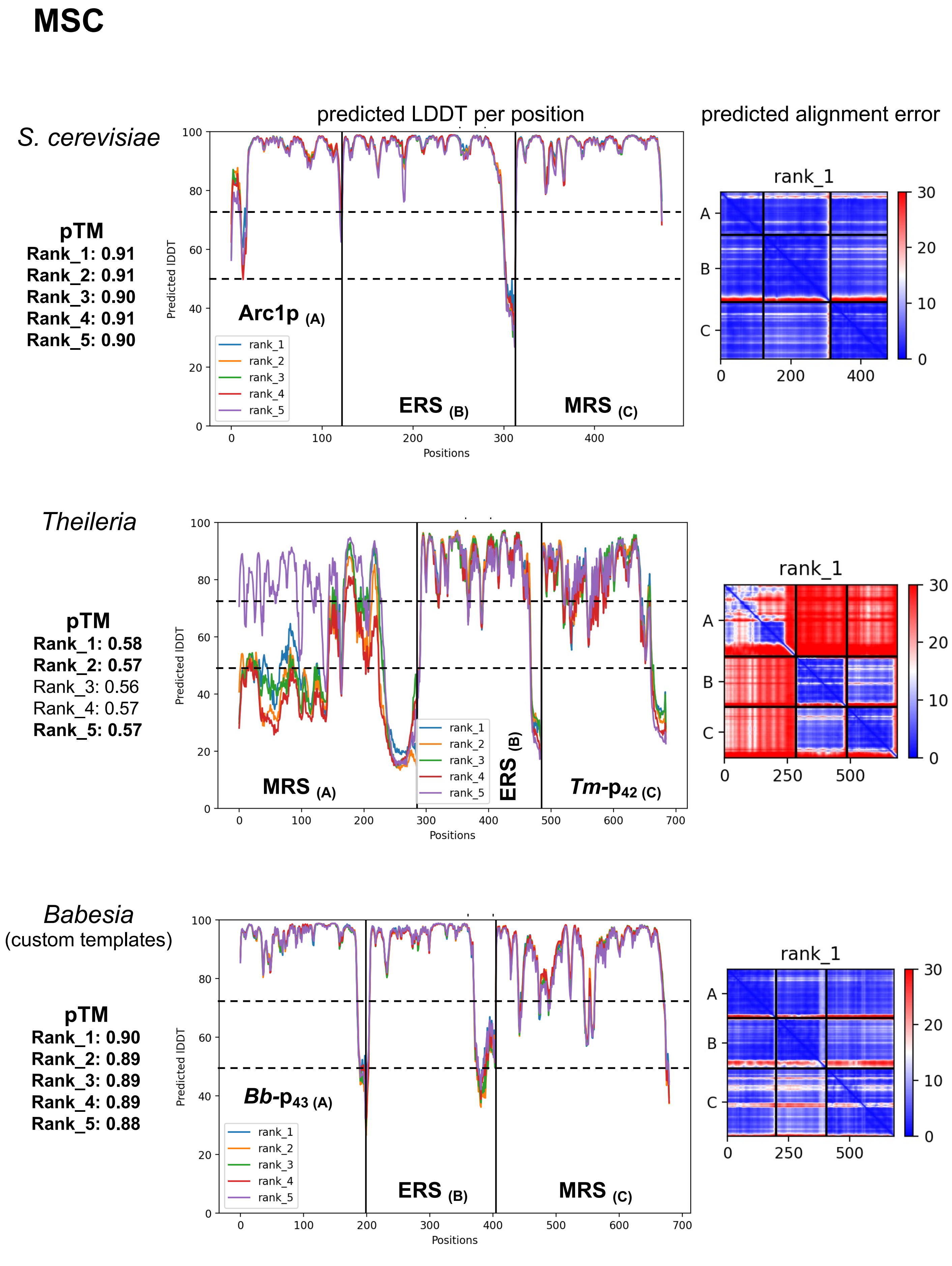
MSC
S. cerevisiae
pTM
Rank_1: 0.91
Rank_2: 0.91
Rank_3: 0.90
Rank_4: 0.91
Rank_5: 0.90
predicted LDDT per position
Arc1p (A)
ERS (B)
MRS (C)
rank_1
rank_2
rank_3
rank_4
rank_5
Predicted lDDT
Positions
predicted alignment error
rank_1
Theileria
pTM
Rank_1: 0.58
Rank_2: 0.57
Rank_3: 0.56
Rank_4: 0.57
Rank_5: 0.57
MRS (A)
ERS (B)
Tm-p42 (C)
Babesia
(custom templates)
pTM
Rank_1: 0.90
Rank_2: 0.89
Rank_3: 0.89
Rank_4: 0.89
Rank_5: 0.88
Bb-p43 (A)
ERS (B)
MRS (C)

**Figure S6**

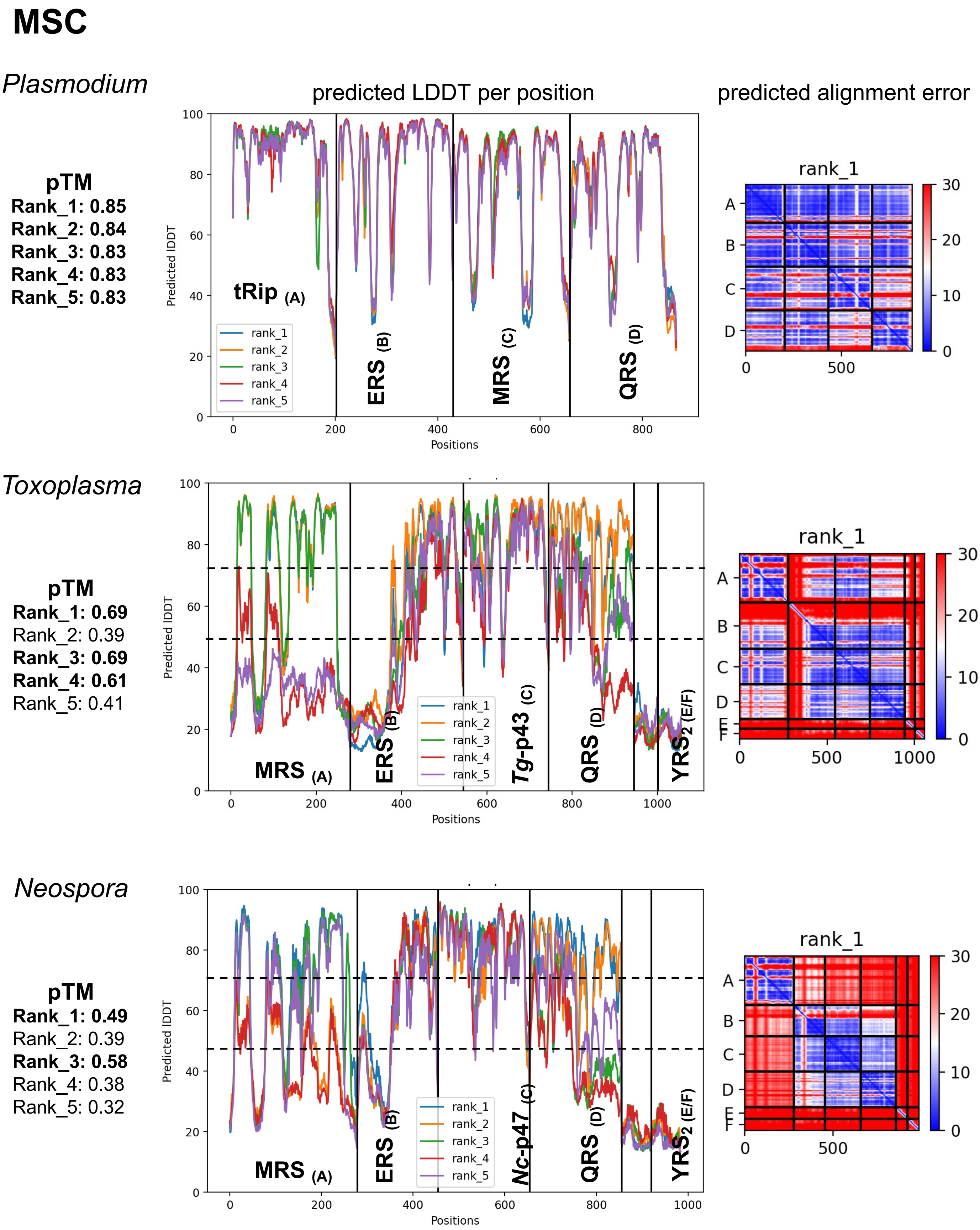


**Figure S6. Confidence metrics of the ColabFold predictions for GST-like interaction network of *S. cerevisiae* and *Apicomplexa* parasites.** See legend Figure S5

### MSC

predicted LDDT per position | predicted alignment error

*Plasmodium*
Q-complex

**pTM**
**Rank_1: 0.70**
Rank_2: 0.66
Rank_3: 0.66
Rank_4: 0.45
Rank_5: 0.43

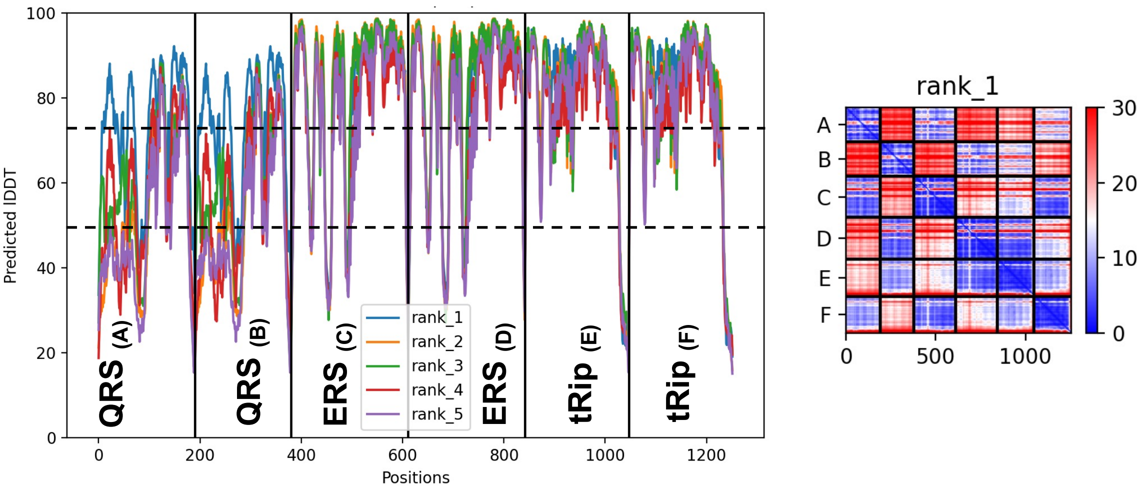


*Plasmodium*
M-complex

**pTM**
**Rank_1: 0.74**
**Rank_2: 0.72**
Rank_3: 0.45
Rank_4: 0.36
Rank_5: 0.36

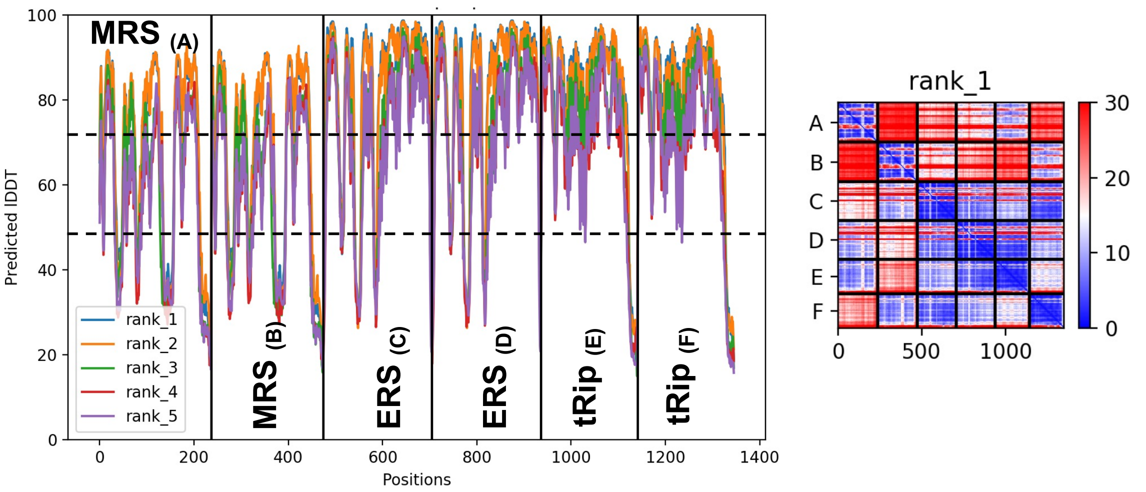


**Figure S7**

### MSC

*Toxoplasma*
Q-complex

**pTM**
**Rank_1: 0.73**
**Rank_2: 0.72**
**Rank_3: 0.66**
**Rank_4: 0.65**
Rank_5: 0.32

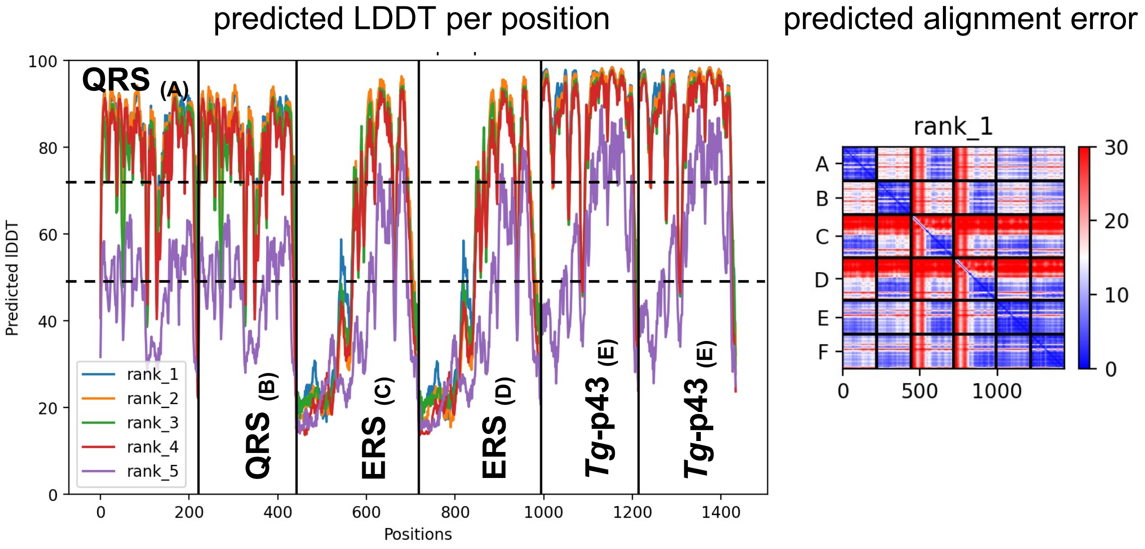


*Toxoplasma*
M-complex

**pTM**
**Rank_1: 0.73**
**Rank_2: 0.69**
**Rank_3: 0.69**
**Rank_4: 0.56**
Rank_5: 0.49

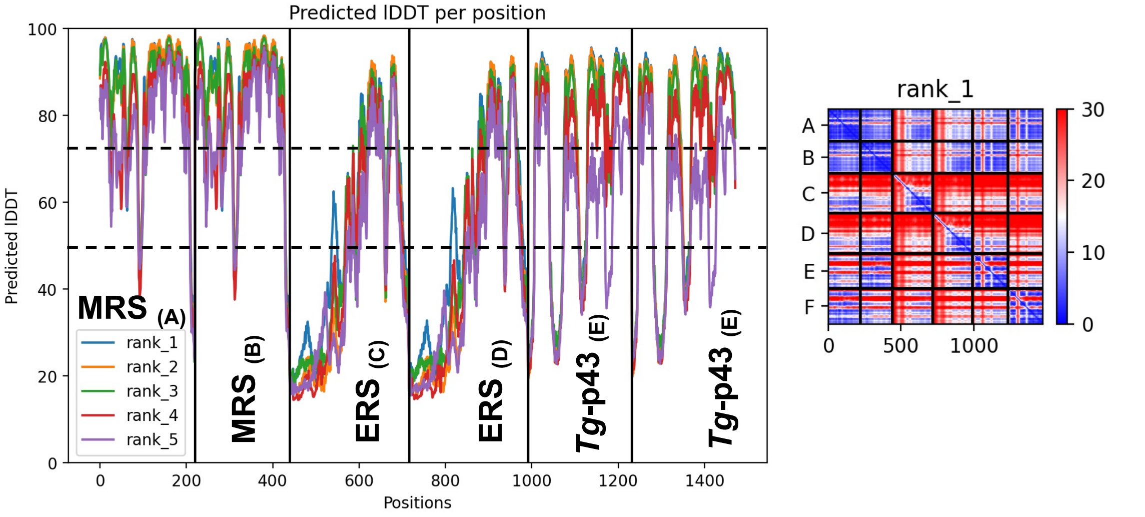


**Figure S7. Confidence metrics of the ColabFold predictions for M- and Q-MSC GST-like backbones in *Plasmodium*, *Toxoplasma* and *Neosporidium*.** See legend Figure S5
